# Direct therapeutic targeting of SWI/SNF induces epigenetic reprogramming and durable tumor regression in rhabdoid tumor

**DOI:** 10.1101/861484

**Authors:** Maggie H. Chasse, Benjamin K. Johnson, Elissa A. Boguslawski, Katie M. Sorensen, Lyong Heo, Zachary B. Madaj, Ian Beddows, Gabrielle E. Foxa, Susan M. Kitchen-Goosen, Bart O. Williams, Timothy J. Triche, Patrick J. Grohar

## Abstract

**Purpose:** Rhabdoid tumor is a pediatric cancer characterized by the biallelic inactivation of SMARCB1, a subunit of the SWI/SNF chromatin remodeling complex. SMARCB1 inactivation leads to SWI/SNF redistribution to favor a proliferative dedifferentiated cellular state. Although this deletion is the known oncogenic driver, SWI/SNF therapeutic targeting remains a challenge.

**Experimental Design:** We define a novel epigenetic mechanism for mithramycin using biochemical fractionation, chromatin immunoprecipitation sequencing (ChIP-seq), and a dual spike-in assay for transposase accessible chromatin sequencing (ATAC-seq). We correlate epigenetic reprogramming with changes with chromatin A/B compartments and promoter accessibility with chromHMM models and RNA-seq. Finally, we demonstrate durable, marked tumor response in an intramuscular rhabdoid tumor xenograft model.

**Results:** Here we show mithramycin and a second-generation analogue EC8042 evict mutated SWI/SNF from chromatin and are effective in rhabdoid tumor. SWI/SNF blockade triggers chromatin compartment remodeling and promoter reprogramming leading to differentiation and amplification of H3K27me3, the catalytic mark of PRC2. Treatment of rhabdoid rumor xenografts with EC8042 leads to marked, durable tumor regression and differentiation of the tumor tissue into benign mesenchymal tissue, including de novo bone formation.

**Conclusion:** Overall, this study identifies a novel therapeutic candidate for rhabdoid tumor and an approach that may be applicable to the 20% of cancers characterized by mutated SWI/SNF.

**STATEMENT OF TRANSLATIONAL RELEVANCE:** There is a tremendous need for novel therapeutic approaches for rhabdoid tumor and the more than 20% of human cancers characterized by dysregulation of the SWI/SNF chromatin remodeling complex. While approaches to target associated complexes, such as PRC2, known to be influenced by dysregulated SWI/SNF are currently being evaluated in the clinic, the direct therapeutic targeting of SWI/SNF has not been explored. Here we identify an inhibitor of SWI/SNF and thoroughly explore the therapeutic development of this compound from a mechanistic and translational perspective thus providing insight into the targeting of this complex as well as a dose, schedule, and biomarker of target inhibition that is immediately clinically translatable.

## INTRODUCTION

The SWI/SNF chromatin remodeling complex is mutated in approximately 20% of human cancers. Biallelic inactivation of SMARCB1, and less commonly SMARCA4, is diagnostic of rhabdoid tumor (1). Rhabdoid tumor can arise in the CNS, kidney, or soft-tissues and is characterized by a poor overall survival of 20-30% (2, 3).

SMARCB1 inactivation does not affect SWI/SNF structural integrity but rather destabilizes the SWI/SNF complex on chromatin (4). This leads to redistribution of SWI/SNF away from promoters and typical enhancers to occupancy of super enhancers (5). SWI/SNF occupancy at super enhancers promotes oncogenesis, drives proliferation, and blocks differentiation (6, 7). Importantly, recent studies have defined a non-canonical SWI/SNF complex as a synthetic lethal target in rhabdoid tumor (8, 9). In addition, the weakened affinity of SWI/SNF for chromatin disrupts antagonism with polycomb repressive complexes (PRC) (10, 11). Due to the antagonistic relationship of SWI/SNF and PRC2, EZH2 small molecule inhibition has been proposed as a promising clinical candidate (12). However, a complementary approach would be to directly target SWI/SNF to reverse the proliferative potential of the cells and perhaps restore the differentiation program.

In this study, we show that the small molecule mithramycin and its second-generation analogue EC8042 inhibit residual SWI/SNF activity. We previously identified mithramycin as an important therapeutic agent in Ewing sarcoma, a tumor known to have dysregulated SWI/SNF (13, 14). In this report, we sought to determine if SWI/SNF dysregulation was a molecular feature contributing to the heightened sensitivity of tumors to mithramycin. We analyzed two independently generated datasets and found that cell lines with mutated or dysregulated SWI/SNF were indeed particularly sensitive to mithramycin and EC8042 (15). Using rhabdoid tumor as a model of dysregulated SWI/SNF, we show that mithramycin directly targets SWI/SNF by evicting the complex from chromatin. Inhibition of residual SWI/SNF activity triggers epigenetic reprogramming that favors promoters and A/B compartment remodeling of specific chromosomes. This reprogramming restores the differentiation program leading to differentiation of rhabdoid tumor cells into benign mesenchymal tissue both in vitro and in vivo. The net effect is a striking durable regression of rhabdoid tumor xenografts and complete cures of a subset of mice treated with just one 3-day infusion of EC8042. Because of the favorable toxicity profile of EC8042, these results establish EC8042 as a novel clinical candidate for rhabdoid tumor and tumors with mutated or dysregulated SWI/SNF.

## MATERIALS & METHODS

### Cell Culture

BT12 and CHLA266 cell lines were obtained from the Children’s Oncology Group, G401 cells from ATCC, TC32 cells from Dr. Lee Helman (Children’s Hospital of Los Angeles) and U2OS cells from Dr. Chand Khanna (Ethos Veterinary Health LLC). Cell lines were routinely monitored for pathogens and cultured as described (16).

### Quantitative real-time PCR (RT-qPCR)

BT12 cells were incubated with mithramycin and the RNA was collected, reverse transcribed, and PCR amplified as previously described (16). Expression was calculated with standard ddCT methods relative to GAPDH and solvent controls.

### Cell Proliferation Assays

IC50s were determined by non-linear regression in duplicate for three independent experiments using MTS assay CellTiter96 and confirmed with real-time proliferation assays on the Incucyte Zoom (16).

### Western Blot

BT12 cells were incubated with mithramycin and lysed with 4% lithium dodecyl sulfate (LDS) buffer. Fractionation lysates were collected with cytoplasmic lysis buffer (25 mM HEPES pH 8.0, 50mM KCl, and 05% NP-40) and the nuclei pellet lysed with 4% LDS buffer. 30 μg of protein was resolved and transferred as described (16). The membranes were blocked with 5% milk in TBS-T and probed with Abcam (H3K27me3), EMD Millipore (SP1), and Cell Signaling (SMARCC1, SMARCE1, H3) antibodies.

### Chromatin Fractionation

3 million BT12, G401, or U2OS were incubated with 100nM mithramycin or PBS control for 8 or 18-hours, washed and collected in PBS and fractionated as previously described (16).

### RNA-Sequencing

RNA was extracted and submitted for 1×75bp sequencing. Libraries were prepared from 500ng of total RNA as described (17). Reads were aligned to hg19 using STAR (v2.7.0f) (18). The index was prepared using default parameters except for -- sjdbOverhang 75 and --sjdbGTFfile, where the Gencode v19 annotations were used. Default parameters were used for alignment with the following modifications: -- readFilesCommand zcat --outReadsUnmapped None --quantMode GeneCounts -- outSAMtype BAM SortedByCoordinate. Gene-level transcript quantification was performed using STAR’s built-in quantification algorithm as noted in the modified alignment parameters. Libraries were normalized using trimmed mean of M-values after filtering for low abundance transcripts using the R (v3.6.1) package edgeR (19, 20). Differential expression analysis was carried out using limma-voom (v3.40.6) (21, 22). Significant genes were determined using a cutoff of q < 0.05. Heatmaps were generated using the pheatmap (v1.0.12) package. GSEA was performed with functional gsea (v1.10.1) package (23). Reactome pathway analysis was performed with the reactomePA (v1.28.0) package (24). Primary explant normal skull osteoblast raw RNA-seq data was downloaded from the SRA study SRP038863 (25). Data were aligned and pre-processed as described above. PCA was carried out using log2(TPM + 1) counts and the prcomp function from the stats package (v3.6.1). Results were plotted using ggplot2 (v3.2.1) and the viridis (v0.5.1) packages.

### Chromatin Immunoprecipitation with high throughput sequencing (ChIP-seq)

BT12 cells were incubated with 100nM mithramycin or PBS control for 8-hours or 18- hours. Cells were cross-linked, lysed, and sheared as described (16). 10µg solubilized chromatin was immunoprecipitated with 1µg mouse IgG and 1µg H3K27me3 (Abcam); 2µg rabbit IgG and 2µg SMARCC1 (Cell Signaling); 1µg rabbit IgG and 1µg H3K27ac (Active Motif). Antibody-chromatin complexes were immunoprecipitated and purified as described (16). ChIP DNA was quantified with SYBR green relative to a standard curve generated with chromatin from the respective sample for each primer set. qPCR as described above was performed with the following primer sets (GAPDH, MYT1, SOX2, CCND1, SP1). Libraries for Input and IP samples were prepared from 10ng of input material and either 10ng or all available IP as described (16).

### ChIP-seq Analysis

Reads were aligned using bwa mem, duplicate marked with samblaster, and filtered and converted to BAM format using samtools (26, 27). BAMs were ingested into R (v3.6.1) and processed using csaw (v1.18.0). A 150bp sliding window with a 50bp step-size was used to summarize the read counts with a maximum fragment size set to 800bp (28, 29). Background was estimated using a 5kb sliding window where reads were binned and summarized and known hg19 blacklist regions were excluded (30). Regions having signal greater than log2(3) fold-change over background were retained for differential analysis. The first principal component was regressed out of the data due to a batch effect being present prior to downstream analysis. Differential analysis was carried out using csaw and edgeR, fitting a quasi-likelihood (QL) negative binomial generalized log-linear model that estimates the prior QL dispersion distribution robustly. Differentially bound regions were generated using an anova-like test or individual contrasts. p-values were combined across clustered sites using Simes’ method to control the cluster false discovery rate as implemented in the combineTests function in csaw. Clustered DBRs were defined as having a q <0.05.

### Assay for Transposase Accessible Chromatin with High Throughput Sequencing (ATAC-seq)

BT12 cells were treated with 100nM mithramycin for 8 or 18-hours or PBS control for 18-hours. 25,000 cells were used to perform omni-ATAC with minor modifications (31). Prior to transposition, 0.1ng lambda phage (Thermo Fisher Scientific) was added to the nuclei pellet. Transposition was carried out for 60-minutes at 37°C. After purification, 0.1ng phiX DNA were added to the transposition DNA prior library amplification. Libraries were amplified and purified as described. Finished libraries were size-selected to retain fragments between 200-800bp using double sided SPRI selection with Kapa Pure Beads. Indexed libraries were pooled and 75bp, paired end sequencing was performed on an Illumina NextSeq 500 sequencer using a 150bp HO sequencing kit (v2). Base calling was done by Illumina NextSeq Control Software (NCS) v2.0 and output of NCS was demultiplexed and converted to FastQ format with Illumina Bcl2fastq2 v2.20.0.

### ATAC-seq Analysis

Reads were aligned to hg19 as described above for ChIP-seq and the enterobacteria phage lambda genome (NC_001416.1) was added as additional contigs. BAMs were processed using csaw (v1.18.0) in R (v3.6.1). A 150bp sliding window with a 50bp step-size was used to summarize the read counts with a maximum fragment size of 500bp. Background was estimated using a 1kb sliding window where reads were binned and summarized. Regions having signal greater than log2(3) fold-change over background were retained for differential analysis. Libraries were normalized using RUVg from the RUVSeq (v1.18.0) package, with the lambda reads as the control “genes” (32). PCA plots examining the effects of increasing k were plotted using EDASeq (v2.18.0) (33). Differentially accessible regions (DAR) were computed by fitting a similar model as described for ChIP-seq, but with the RUV weights added. Statistical significance was determined as q < 0.05. Library complexity analysis was performed using preseqR (v4.0.0) (34). Chromatin conformation was inferred by extending methods as previously described and implemented in compartmap (v1.3). Filtered read counts are summarized within a bin, pairwise Pearson correlations are computed across samples within a group, and the first principle component describes the chromatin conformation state using the sign of the eigenvalue. Chromosome-wide compartment dissimilarity scores were computed relative to solvent by calculating 1 - Pearson correlations.

### chromHMM Analysis

ChIP-seq data was downloaded from phs000470 for 19 MRT patients and aligned to the hg38 assembly with bwa mem, duplicates marked and removed with samblaster, and converted to BAM format with samtools (35). The following chromatin marks were used as input to construct the chromHMM (v1.18) model: H3K4me1, H3K4me3, H3K9me3, H3K27Ac, H3K27me3, H3K36me3 (36, 37). Additionally, matched input samples were used for local thresholding during the binarization step. BAMs were binarized using default values and segmented using the Roadmap 18-state core model collapsed into 6 “super-states” (38). The original and collapsed chromatin state calls were combined into a RaggedExperiment (v1.8.0) object. Differential regions were queried for overlaps to specific states, where a consensus state was called as having at least 50% of patient samples having the same inferred state from chromHMM. Donut plots were generated using significant (q < 0.05) trended changes in accessibility (ATAC-seq) or binding (ChIP-seq) from 8 to 18 hours of mithramycin treatment relative to solvent.

### siRNA Knockdown

RNAiMax Lipofectamine was added to siRNA targeting SMARCA4, SMARCC1, or SP1 and allowed to complex. BT12 cells were added to the mixture and incubated for 30h (SP1) or 48h (SMARCA4 and SMARCC1) before collection for qRT-PCR analysis.

### Luciferase Cells

CMV-luciferase plasmid was linearized with HF-Sal1 and transfected into G401 cells as described (13). Cells were expanded under G418 selection and confirmed to be pathogen-free.

### Xenograft Experiments

5×10^6^ G401-luc cells were injected intramuscularly into the gastrocnemius of 8-10 week old female homozygous nude mice (Crl; Nu-*Foxn1^Nu^*). Tumors were established to a minimum of 100 mm^3^ and tumor volume was measured daily by caliper and the volume determined as described (16). Mice (n=12 per cohort) were treated with mithramycin given at 1mg/kg mithramycin IP (8mg/kg total), 2.4mg/kg in a 3-day or 7-day continuous infusion alzet micro-osmotic pump (model 1007D) or vehicle (PBS supplemented with magnesium or calcium) per the identical schedules (n =12). EC8042 treated mice were treated with 30mg/kg in a 3-day infusion or 50 mg/kg in a 7-day infusion. All experiments were performed in accordance with and the approval of the Van Andel Institute Institutional Animal Care and Use Committee (IACUC). Investigators were not blinded to the treatment groups.

### Bioluminescence Imaging

Mice were administered Firefly D-Luciferin with intraperitoneal injections. After injection anesthesia was administered throughout the image acquisition (3% isoflurane at 1L/min O2 flow). Bioluminescent images were taken 10 minutes after injection using the AMI- 1000 imaging system.

### Microcomputed Tomography (micro-CT)

Mineralized tissue within tumors was examined using the SkyScan 1172 micro-computed tomography system (Bruker MicroCT). Tumors were scanned in 70% ethanol using an X-ray voltage of 60kV, current of 167µA, and 0.5mm aluminum filter. The pixel resolution was set to 2000×1200, with an image pixel size of 8µm. A rotation step of 0.40 degrees and 360° scanning was used. 2D cross-sectional images were reconstructed using NRecon 1.7.4.6. A volume of interest (VOI) was defined for each tumor using DataViewer 1.5.6.3. A region of interest (ROI) around the tumor was defined and 3D files were generated using CTAn 1.18.8.0. Representative 3D images were created using CTvol 2.3.2.0.

### Tissue Staining and Immunohistochemistry

Tissues were decalcified in 10% EDTA (pH 8.0) and sectioned following paraffin embedding as described (16). Tissue was incubated with H3K27me3 (Abcam, 1:250), Cleaved Caspase-3 (Cell Signaling, 1:250), H3K27ac (Abcam, 1:1250), or human mitochondria (Abcam, 1:800) washed and then secondary antibody (Envision+System HRP labelled polymer Anti-Rabbit, Dako 1:100).

### Project Statistics

qPCR data is normalized to solvent (mRNA expression data) or input (ChIP data) as fold-change from 3 independent experiments performed in technical duplicate. The p-values were determined by one-way ANOVA using Dunnet test for multiple comparisons.

### Data Availability

The sequencing data has been deposited on GEO under the accession number GSE137404.

## RESULTS

### Mithramycin sensitivity is linked with SWI/SNF mutation or complex dysregulation

In order to determine if dysregulated SWI/SNF is associated with heightened sensitivity of cells to mithramycin, we analyzed two independently published data sets (Figure 1A, S1A) (15, 39). Indeed, we found that 19 of the top 25 most sensitive sarcoma cell lines have SWI/SNF mutation or dysregulation (Figure 1B). In addition, rhabdoid tumor cell lines which are known to be driven only by SMARCB1 deletion exhibited a striking sensitivity to mithramycin that paralleled Ewing sarcoma. Mithramycin is known to target the oncogenic driver of Ewing sarcoma, EWS-FLI1, and dysregulated SWI/SNF is a hallmark of both tumors. Therefore, it follows that this heightened sensitivity may be related to SWI/SNF.

**Figure 1:**
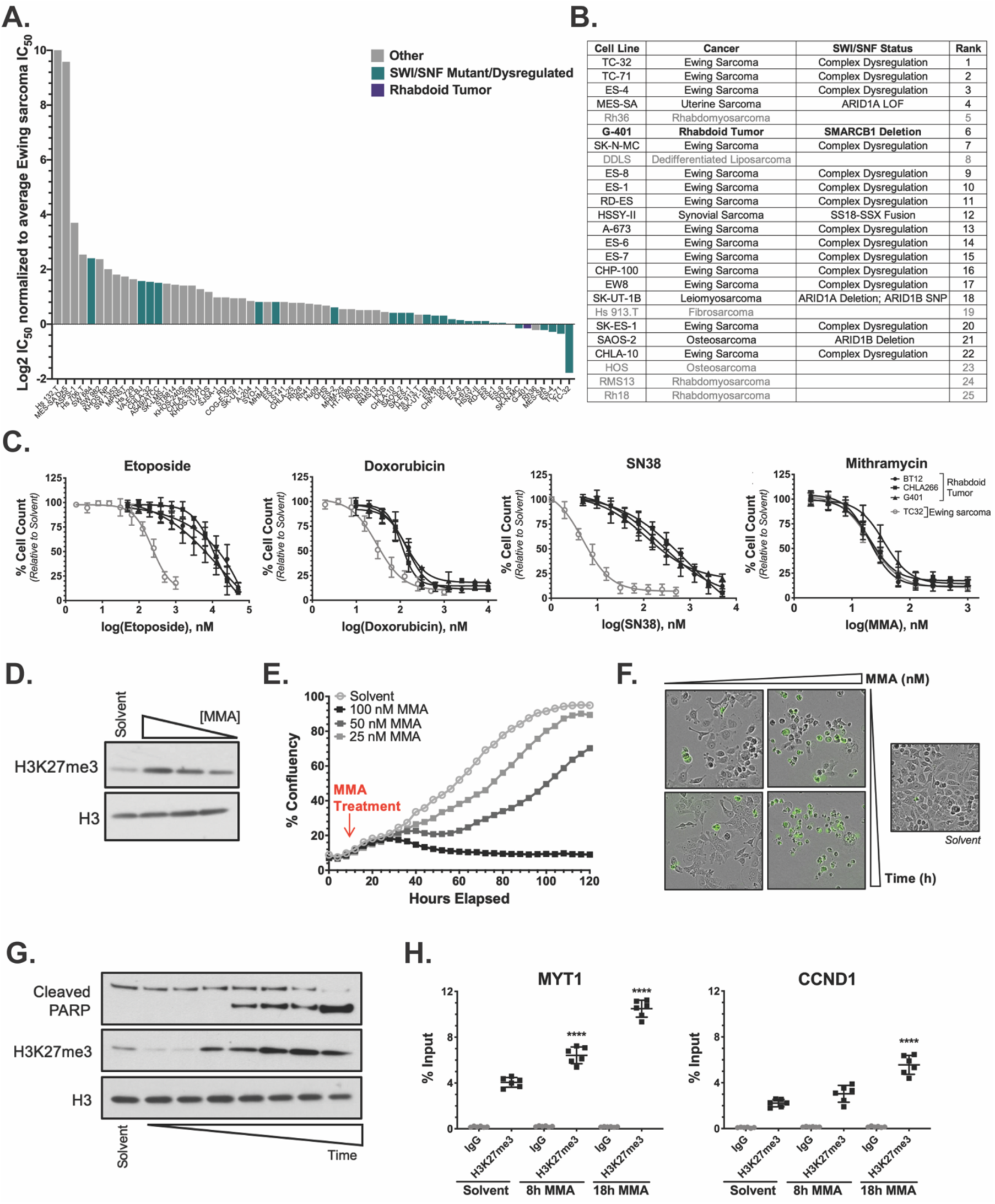
Mithramycin cellular sensitivity favors SWI/SNF mutant cancers. **A:** Graph of IC50 as a function of cell line generated from a published screen of 445 agents in 63 sarcoma cell lines (39). Cell lines with mutated or dysregulated SWI/SNF (green) cluster towards the right on the graph indicating these cell lines are more sensitive to mithramycin. **B:** Table highlighting the SWI/SNF dysregulation status in the top 25 sarcoma cell lines from Fig. 1A. Mutation status was confirmed in COSMIC (49) or the DepMap (https://depmap.org/portal/). **C:** Dose response curves of rhabdoid tumor and Ewing sarcoma cell lines. RT cell lines (black) are sensitive to mithramycin treatment with a similar IC50 value as TC32 ES cells (grey). RT cell lines are not sensitive to three broadly-active chemotherapeutic agents: etoposide, doxorubicin or SN38. **E:** Western blot showing concentration-dependent increase in H3K27me3 following exposure to 100nM, 50nM, 25nM mithramycin for 18h in BT12 cells relative to loading control (H3). **E:** Mithramycin induces a concentration-dependent suppression of proliferation in BT12 cells. Cells were exposed to 25nM, 50nM, or 100nM MMA for 18h, replaced with drug- free media and monitored using live cell imaging. **F:** Mithramycin induces morphological changes and apoptosis relative to control cells (solvent). BT12 cells treated with 25nM (left) or 100nM (right) mithramycin for 18h (top) or 48-hours (bottom) in the presence of cleaved caspase 3/7 reagent that fluoresces with caspase activation. **G:** Mithramycin leads to H3K27me3 amplification in a time-dependent manner that precedes the induction of apoptosis as measured by the cleavage of PARP. Western blot lysates collected at 1h, 2h, 4h, 8h, 12h 16h, 18h of continuous 100nM mithramycin treatment. **H:** Chromatin immunoprecipitation qPCR (ChIP-qPCR) of H3K27me3 at *MYT1* and *CCND1*. H3K27me3 occupancy is increased in a time-dependent manner.

In order to confirm that this sensitivity is related to common molecular features of these tumors and exclude non-specific mechanisms, we compared the cellular sensitivity of these tumors to standard chemotherapeutic agents. Three RT cell lines (BT12, CHLA266, G401) were 70-fold, 4-fold, and 100-fold less sensitive to three broadly active chemotherapy agents, etoposide, doxorubicin, and SN38 (the active metabolite of irinotecan) than Ewing sarcoma cells (Figure 1C). In contrast, we confirmed that all three cell lines were equally sensitive to mithramycin as the Ewing sarcoma cells. These data exclude a general sensitivity of RT cells to non-specific DNA damaging agents as rhabdoid tumor is known to be chemo-resistant. In addition, these findings are consistent with a striking sensitivity of rhabdoid tumor to mithramycin that is associated with SWI/SNF dysregulation.

### Mithramycin evicts SWI/SNF from chromatin and amplifies H3K27me3

Due to the known effects of mithramycin in Ewing sarcoma cells on EZH2, we reasoned that MMA may be acting to inhibit EZH2. However, mithramycin treatment led to an impressive concentration-dependent increase in H3K27me3 (Figure 1D). We correlated H3K27me3 amplification with suppression of cellular proliferation and induction of apoptosis as measured by cleaved caspase (Figure 1E, 1F). Further, treatment with mithramycin led to time-dependent H3K27me3 amplification that correlated with cleavage of PARP (Figure 1G). We validated H3K27me3 amplification at specific loci associated with SWI/SNF with chromatin immunoprecipitation. In a time-dependent manner, H3K27me3 occupancy increases at *MYT1* and *CCND1* with no change of occupancy at a locus control, *GAPDH* (Figure 1H, S1B). These data are consistent with a further reduction of SWI/SNF activity at these loci leading to increased PRC2 activity.

In order to test this hypothesis, we treated RT cells with 100nM mithramycin and biochemically fractionated the cells into chromatin or soluble fractions. Mithramycin evicts SWI/SNF from chromatin by 8-hours of exposure (Figure 2A). This effect also occurred in G401 RT cells but was not seen in U2OS, a wildtype SWI/SNF osteosarcoma cell line, suggesting some selectivity of this effect for mutant SWI/SNF (Figure 2B, 2C). It is notable that both mithramycin and SWI/SNF bind the DNA minor groove and therefore mithramycin may be competitively evicting SWI/SNF as has been suggested with a structurally related compound, chromomycin (40, 41). We confirmed loss of SWI/SNF from chromatin with ChIP-qPCR of SMARCC1 where SMARCC1 occupancy decreases at *MYT1* and *CCND1* relative to *GAPDH* (Figure 2D, S1B).

**Figure 2:**
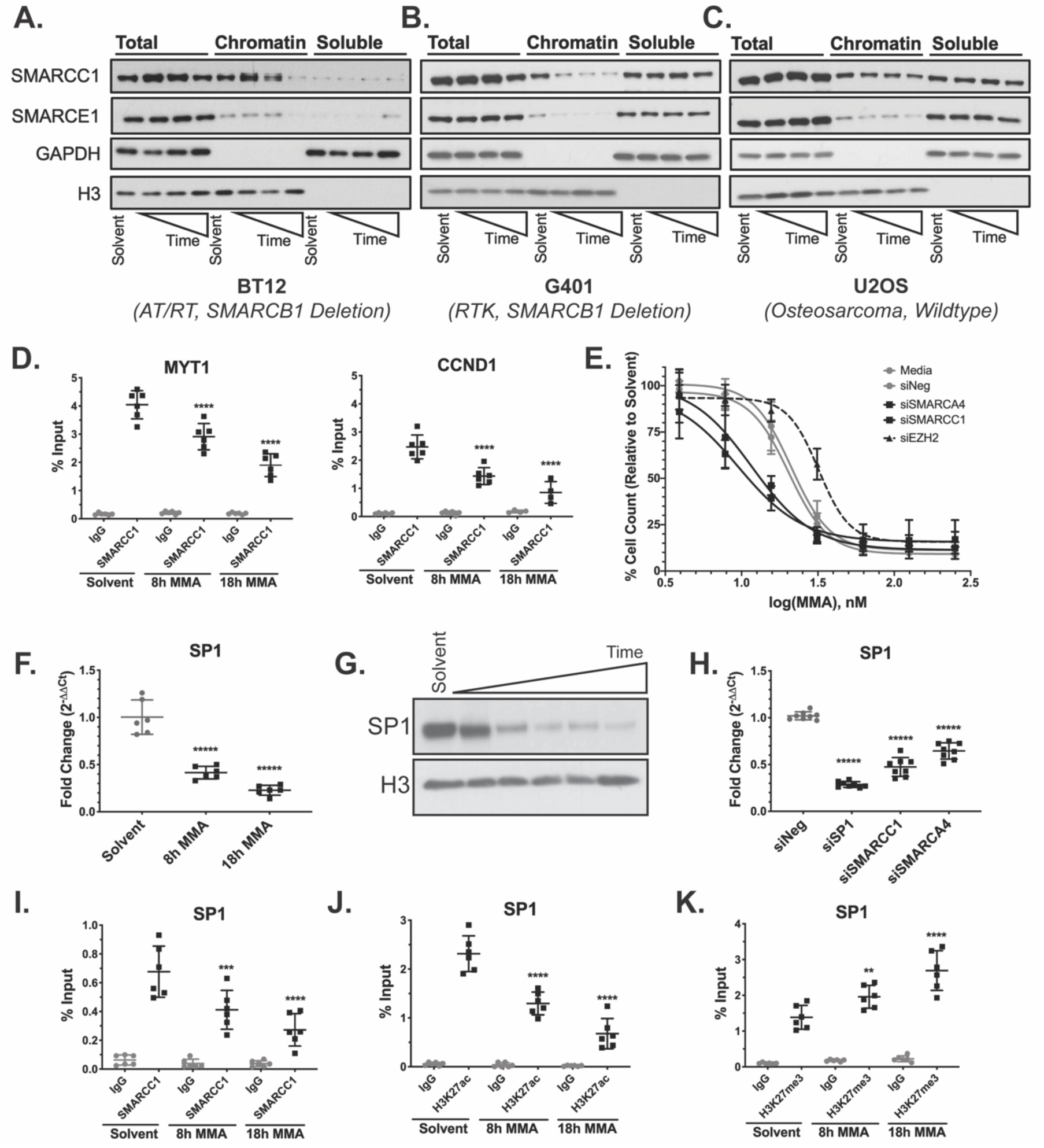
Mithramycin induced morphological changes are dependent on SWI/SNF eviction and the induction of H3K27me3. **A, B, C:** Mithramycin evicts SMARCC1 and SMARCE1 SWI/SNF subunits from chromatin in a time-dependent manner in BT12 **(A)** and G401 **(B)** cells but not U20S **(C)** cells. Western blot analysis showing whole cell lysate (total), chromatin bound (chromatin), and nuclear and cytoplasmic soluble (soluble) fractions collected after exposure to 100nM mithramycin for 1, 8 or 18h and probed for the SWI/SNF subunits (SMARCC1 or SMARCE1) or H3 (chromatin control) and GAPDH (soluble control). **D:** Loss of SWI/SNF occupancy at defined loci in the genome as measure by ChIP- qPCR at known SWI/SNF target genes, *MYT1* and *CCND1*. **E:** Suppression of SWI/SNF subunit expression sensitizes BT12 cells to mithramycin while suppression of EZH2 antagonizes mithramycin activity. Data represents dose response curves of mithramycin in BT12 cells following a 48h exposure in the presence of siRNA silencing of SMARCA4 or SMARCC1 SWI/SNF subunits relative to untreated cells (media), a non-targeting siRNA (siNeg) or the PRC2 subunit EZH2. **F:** SP1 mRNA expression is reduced following 100nM mithramycin treatment as measured by qPCR fold-change relative to GAPDH (2^ddCT^). ****, p-value <0.0001. **G:** SP1 protein expression tracks with the reduction in mRNA. Western blot showing suppression of SP1 expression compared to loading control (H3) after 100nM mithramycin exposure for 1h, 4h, 8h, 12h, 18h. **H:** SP1 expression is dependent on SWI/SNF. siRNA silencing of SMARCC1 and SMARCA4 subunits is associated with a similar loss of SP1 expression as direct silencing of SP1 as measured by qPCR relative to GAPDH (2^ddCT^). ****, p-value <0.0001. **I-K:** The loss of SP1 expression is associated with a decrease in SWI/SNF occupancy of the SP1 promoter and a change from H3K27ac to H3K27me3. Data represents ChIP- qPCR analysis following 8-hours or 18-hours of 100nM mithramycin exposure and immunoprecipitation of SMARCC1 **(I)**, H3K27ac **(J)**, and H3K27me3 **(K)**. ****, p-value <0.0001; ***, p-value <0.001; **, p-value <0.01.

Finally, we confirmed the relationship of SWI/SNF and PRC2 to mithramycin sensitivity using siRNA to silence SMARCA4 and SMARCC1 as well as EZH2. Further reduction of SWI/SNF sensitizes BT12 cells to mithramycin while EZH2 loss confers resistance (Figure 2E). These data provide further evidence that the mechanism of growth suppression is SWI/SNF inhibition triggering PRC2 activation leading to increased H3K27me3 and reversal of the oncogenic program.

### Mithramycin evicts SWI/SNF from the SP1 promoter decreasing expression

Mithramycin is widely regarded as a SP1 inhibitor (42, 43). To exclude SP1 blockade as the mechanism of growth impairment, we looked at the impact of mithramycin on SP1. We found that mithramycin decreases SP1 expression in rhabdoid tumor cells consistent with reports in other cell types (Figure 2F, 2G). We reasoned that SWI/SNF eviction may be driving epigenetic reprogramming at the SP1 promoter to account for the observed inhibitory effects on SP1. Consistent with this hypothesis, knockdown of SMARCC1 and SMARCA4 led to a loss of SP1 expression (Figure 2H). In addition, SWI/SNF occupancy did decrease at the SP1 promoter as shown by ChIP-qPCR of SMARCC1 relative to the *GAPDH* control (Figure 2I, S1B). Further, the loss of SWI/SNF binding was associated with increased H3K27me3 and reduced H3K27ac (Figure 2J, 2K). These data indicate the effect of mithramycin on SP1 activity occurs at the SP1 promoter through eviction of SWI/SNF.

In order to determine if SP1 is a vulnerability in RT cells, we treated BT12 and G401 cells with tolfenamic acid, a non-steroidal anti-inflammatory that degrades SP1 protein. However, despite a loss of SP1 expression, RT cells were not sensitive to tolfenamic acid (Figure S1C, S1D). SP1 knockdown with siRNA confirmed that SP1 loss did not impact BT12 cell viability (Figure S1D). Cumulatively, these data indicate SP1 loss is not the primary driver of mithramycin sensitivity in rhabdoid tumor. In addition, in these cells, the mechanism of SP1 suppression is epigenetic occurring by SWI/SNF eviction, H3K27me3 accumulation, and an associated loss of H3K27ac.

### Mithramycin induces epigenetic reprogramming of promoters and chromatin compartments

We next sought to determine if SWI/SNF eviction functions in an analogous fashion at other promoters to drive gene-expression changes responsible for the cellular hypersensitivity to mithramycin. We performed ATAC and ChIP sequencing following mithramycin exposure of BT12 cells. To quantify chromatin remodeling, we developed novel spike-in controls for ATAC-seq where non-chromatinized lambda phage was used for library complexity normalization (Figure S2, S3). In order to correlate accessibility with promoters and enhancers, we performed H3K27ac ChIP-sequencing and normalized the data with the ATAC-seq lambda phage reads.

Recent studies identified a non-canonical SWI/SNF complex as a synthetic lethal RT target that binds promoters and CTCF motifs to drive oncogenic programs (9). This model suggests SWI/SNF eviction would elicit both regional focal changes, likely at promoters, as well as CTCF-dependent chromatin compartment remodeling. To gain insight into chromatin accessibility changes following mithramycin treatment, we quantified the number of regions (q-value <0.05) with differential H3K27ac and accessibility for each chromosome (Figure 3A). Overall, the distribution of H3K27ac and accessible chromatin across the genome was relatively uniform. Differential regions at 8-hours of mithramycin treatment continue to trend at 18-hours and were correlated with changes near the transcriptional start site (Figure 3B, S4).

**Figure 3:**
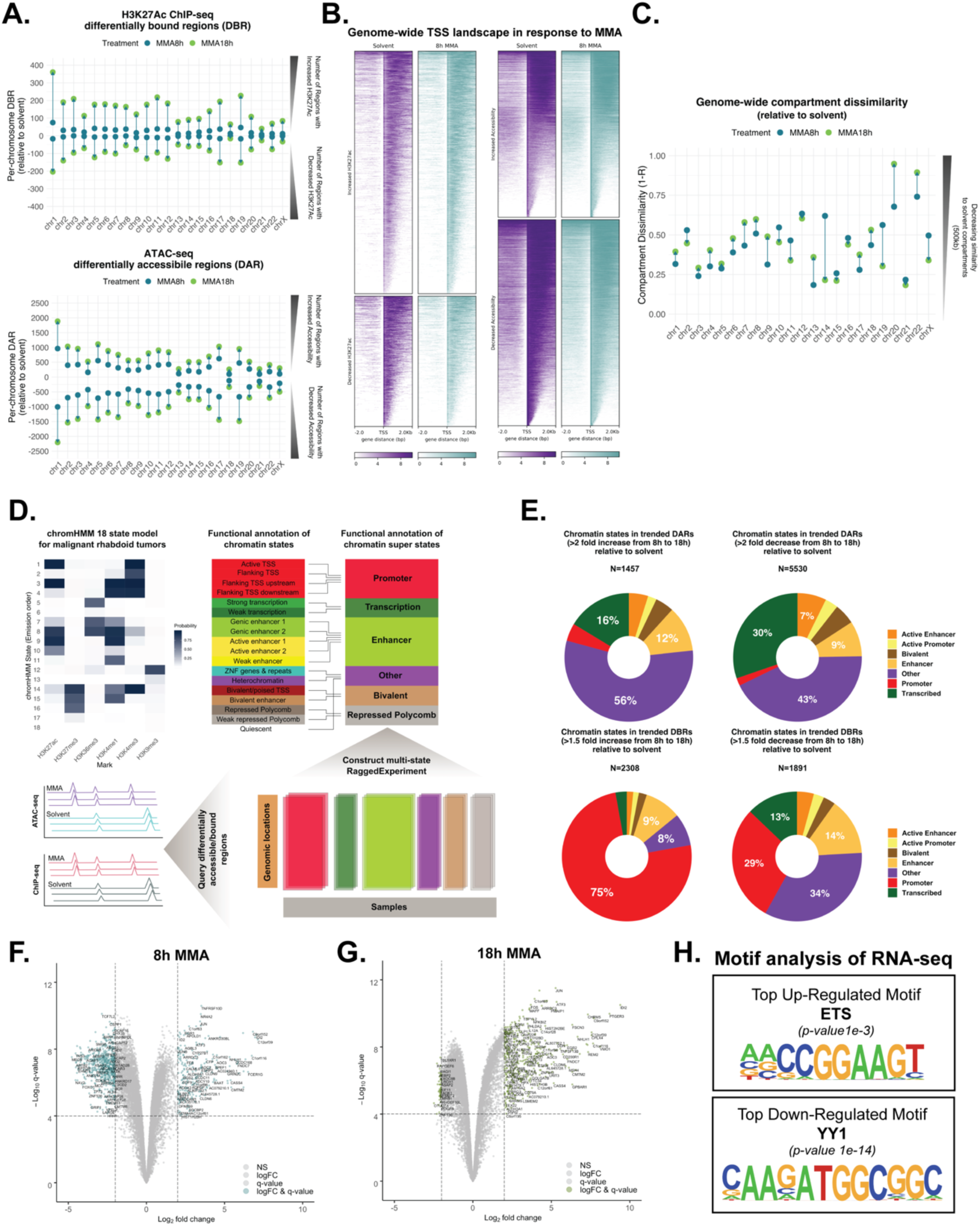
Mithramycin induces epigenetic reprogramming of chromatin compartments and promoters. **A:** Dumbbell plots representing the number of differentially bound (left) or accessible (right) regions per chromosome as determined by chromatin immunoprecipitation of H3K27ac (left) and ATAC-seq (right) following treatment with mithramycin for 8 (blue) or 18h (green). **B:** Heatmaps showing H3K27ac ChIP-seq differentially bound regions (left) and ATAC-seq differentially accessible regions (right) following 8h mithramycin treatment. Peaks were filtered with a 10e-5 q-value threshold. A 2kb window is centered on the TSS. **C:** Dumbbell plots representing ATAC-seq chromatin compartments per chromosome following mithramycin treatment for either 8 (blue) or 18h (green). **D:** Schematic for the 18 state chromHMM model built for MRTs and collapsed into 6 super states. Chromatin states were called if the state was present in at least 50% of samples. ATAC-seq and H3K27ac ChIP-seq peaks were queried against the 6 super states. **E:** Donut plots representing the percentage of each chromatin super state across treatment time (from 8h to 18h) that increased 2-fold (left) or decreased 2-fold (right). **F, G:** Volcano plot showing gene expression trends at 8h **(F)** and 18h **(G)** MMA treatment. Dashed lines represent a 2 logFC and 10e-5 q-value threshold. **H:** Motif analysis of the top up-regulated and down-regulated motifs from the RNA-seq gene lists that pass the logFC and q-value threshold in 4F and 4G.

ATAC-seq motif analysis identified CTCF was the top down-regulated motif. Due to the known relationship of CTCF with topologically associated domains, we explored how mithramycin treatment affects A/B chromatin compartments (Figure 3C). In this case, we found a more dynamic pattern of compartment reorganization where specific chromosomes showed substantial changes compared with the regional changes in Figure 3A. Here, chromosomes trend in dissimilarity to solvent from 8-hours to 18-hours treatment (e.g. chr9) while other chromosomes trend back toward solvent by 18-hours (e.g. chr14). Strikingly, for chromosomes that did not have large region differences (e.g. chr20, 22), these chromosomes exhibited compartment remodeling with mithramycin treatment (Figure 3A, 3C). These data indicate that mithramycin treatment leads to both changes in the number of accessible regions as well as compartment remodeling in the absence of regional accessibility changes consistent with the described model of non-canonical SWI/SNF activity in rhabdoid tumor.

In order to understand how these changes in compartments influence the chromatin states specific to rhabdoid tumor, we developed chromHMM tracks from primary rhabdoid tumors sequenced in TARGET collapsed into six super-states (Figure 3D). We overlaid the differential H3K27ac ChIP-seq and ATAC-seq peaks with the chromHMM super-states. Again, we found evidence of compartment remodeling with the largest percentage of DARs being in heterochromatin (Figure 3E). Due to the established relationship of SWI/SNF to enhancers, we hypothesized that mithramycin would lead to rewiring of the enhancers. However, while 480 of the H3K27ac peaks aligned with enhancer states, 2281 peaks overlaid promoters (184 active or bivalent promoters). These peaks correlated with 75% of the DBRs that increase in the H3K27ac ChIP-seq mapped to the promoters while 30% of promoters decreased. The ATAC-seq peaks were divided among enhancer (661), promoter (172) and transcribed (1912) regions. These data indicate mithramycin primarily remodels RT promoters and gene bodies rather than enhancers.

Promoter reprogramming correlated with RNA-sequencing at 8-hour and 18-hours mithramycin treatment and with the described mechanism of action (Figure 3F, 3G). Volcano plots show an increase in the number of up-regulated genes at 8-hours (210 genes) compared to 18-hours (368 genes). Although a large number of genes decreased in expression (615 genes) at 8-hours, by 18-hours only 49 genes showed a decrease in expression. Gene ontology analysis reflected the described mechanism as chromatin modifying enzymes and chromatin organization pathways were downregulated at 8-hours while PRC2 methylation of histones was enriched in the up- regulated pathways at 18-hours (24). Finally, we correlated gene expression changes with transcription factor consensus motifs (Figure 3H). ETS is the top up-regulated motif which is known to be associated with SWI/SNF (14). YY1 is the top down-regulated motif which is consistent with the idea that mithramycin remodels chromatin compartments to reprogram enhancer-promoter loops (44). These data exclude a general effect on transcription as causative for the cellular sensitivity and instead are consistent with removal of SWI/SNF triggering gene expression changes to drive a specific cellular phenotype.

### SWI/SNF inhibition by mithramycin drives divergent phenotypes in rhabdoid tumor cells

In order to link these chromatin changes that occur with treatment to a defined cellular phenotype, we integrated the genomic analyses to identify gene expression changes that are associated with specific changes in promoter accessibility and activation. We found a complex transcriptional program that favored both apoptosis and cellular differentiation. Analysis of genes that had at least a 1 logFC change at both mithramycin exposures led to 54 common genes up-regulated and 25 common genes down- regulated (Figure 4A, S5C). The most significant enriched processes in the up-regulated genes were related to cell death and apoptosis; an observation also seen with gene set enrichment analysis (Figure 4A, 4B). These transcriptional changes were directly related to promoter reprogramming of, for example, *BCL10*, *BTG2*, and *CDKN1A* (Figure 4C, S5D). We also found striking evidence for differentiation signatures. GSEA identified a stem cell differentiation program and clearly demonstrated reversal of the 10 differentiation gene signatures pathways previously independently identified as aberrantly activated in primary AT/RT (Figure 4D, 4E, 4F).

**Figure 4:**
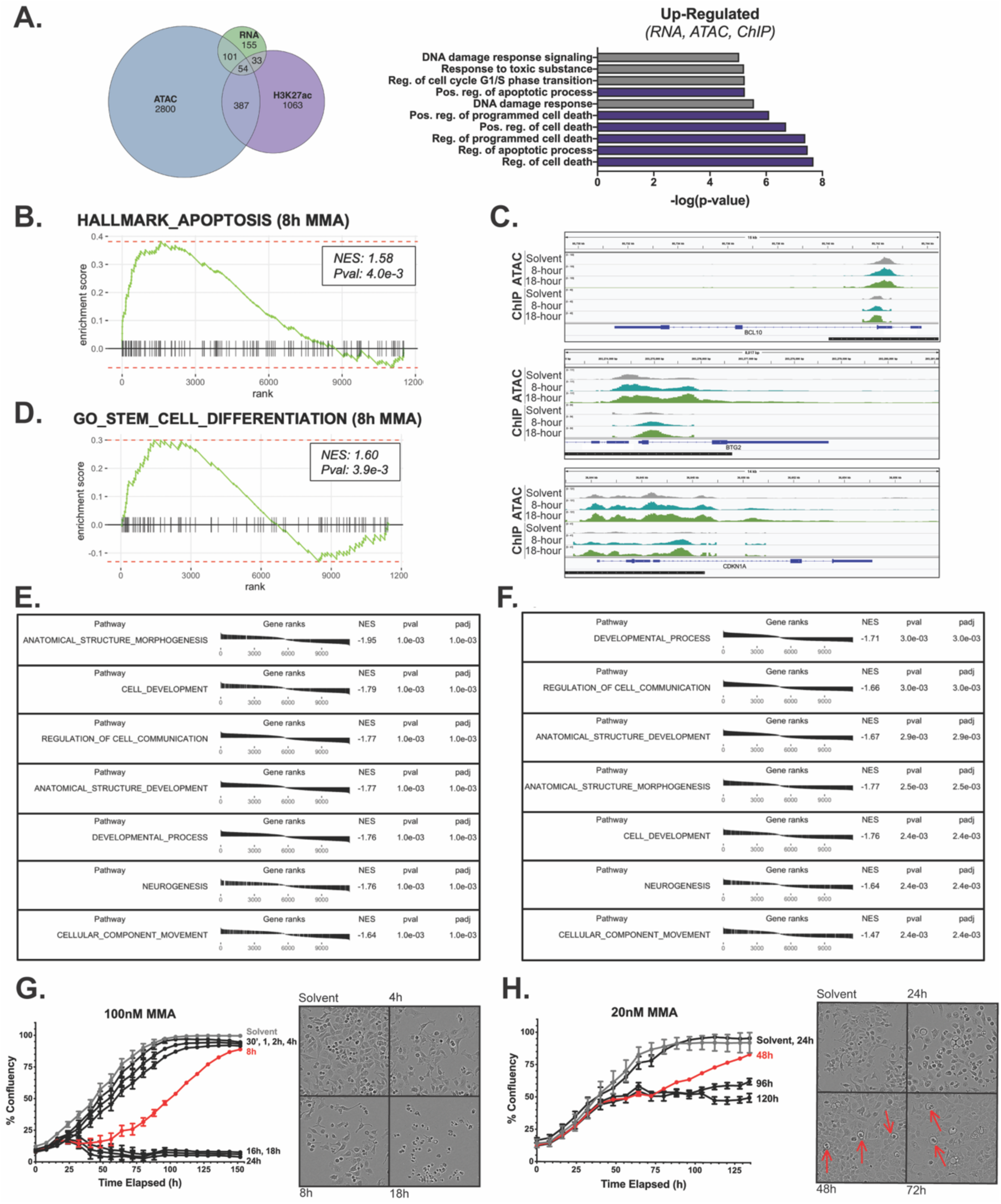
SWI/SNF inhibition by mithramycin drives divergent phenotypes in rhabdoid tumor cells. **A:** Venn diagram showing overlap of the 54 genes that were up-regulated by at least 1- logFC in the RNA-seq, ATAC-seq, and H3K27ac ChIP-seq datasets at both 8h and 18h of mithramycin treatment. 6 of the top 10 pathways up-regulated relate to cell death and apoptosis. **B:** fgsea analysis of the apoptosis hallmark gene set shows enrichment following 8h of mithramycin treatment (p-value 4.0e-3). **C:** IGV tracks of up-regulated genes in the multi-omic analysis (bottom). *BCL10* and *BTG2* play crucial roles in apoptosis while *CDKN1A* is a cell cycle progression gene. Black bars indicate promoters from hg19 within 2kb of the TSS. **D:** fgsea analysis of the stem cell differentiation GO gene set shows enrichment following 8-hours of mithramycin treatment (p-value 3.9e-3). **E, F:** fgsea analysis of gene ontology terms upregulated in primary AT/RT tumors as described in (5) following 8h **(E)** or 18h mithramycin treatment **(F)**. Mithramycin downregulates the expression of these pathways. **G:** BT12 cells show a threshold of exposure that leads to irreversible growth inhibition. The cells were exposed to 100nM MMA for the indicated times followed by replacement with drug-free medium. Beyond 8h (red) of mithramycin exposure, the cells do not recover proliferative potential and exhibit a phenotype consistent with cell death. **H:** BT12 cells treated with 20nM MMA show a similar exposure threshold but different cellular phenotype than with 100nM. The cells were exposed to 20nM MMA for the indicated times followed by replacement with drug-free medium. Beyond 48h of exposure the cells do not recover. There is evidence of mesenchymal differentiation and the appearance of maturing

Importantly, both of the observed transcriptional genotypes were reflected in cellular phenotype. We exposed BT12 and G401 cells to mithramycin for defined time periods and evaluated the cellular morphology using live-cell imaging. BT12 and G401 cells showed evidence of cell death and no longer recovered proliferative potential with 8- hours of 100nM mithramycin (Figure 4G, S5E). At lower concentrations but longer exposure of mithramycin, the cells remained growth arrested and showed evidence of lipid accumulation and mesenchymal differentiation with the appearance of maturing adipocytes (Figure 4H). These data clearly correlate genotype with cellular phenotype and shows that transient high-concentration exposure favors an apoptotic phenotype while continuous exposures to mithramycin favors differentiation.

### Mithramycin shows activity in an intramuscular rhabdoid tumor xenograft

In order to determine which of the exposures is more effective in this tumor type, we directly compared transient high-dose exposure to mithramycin given by intraperitoneal injection to mice bearing RT xenografts to continuous exposure administered by a surgically implanted osmotic diffusion pump. We chose an intramuscular xenograft model with G401 cells to exclude limitations on the CNS penetration of the drug and to model the sarcoma variant of the tumor. In addition, a renal capsular model showed erratic growth (data not shown). Mice with established tumors of a minimum size of 100 mm^3^ were treated with 1mg/kg of mithramycin given by intraperitoneal injection three times per week (every other day) for 2 weeks or 2.4mg/kg given over 3 days with an Alzet pump. Bolus injection showed limited tumor growth suppression (Figure 5A). In contrast, mice treated with a continuous infusion led to a more pronounced suppression of tumor growth, including a regression of a tumor that was 500mm^3^ at the start of treatment, and persisted for almost 2 weeks after discontinuing treatment (Figure 5A).

**Figure 5:**
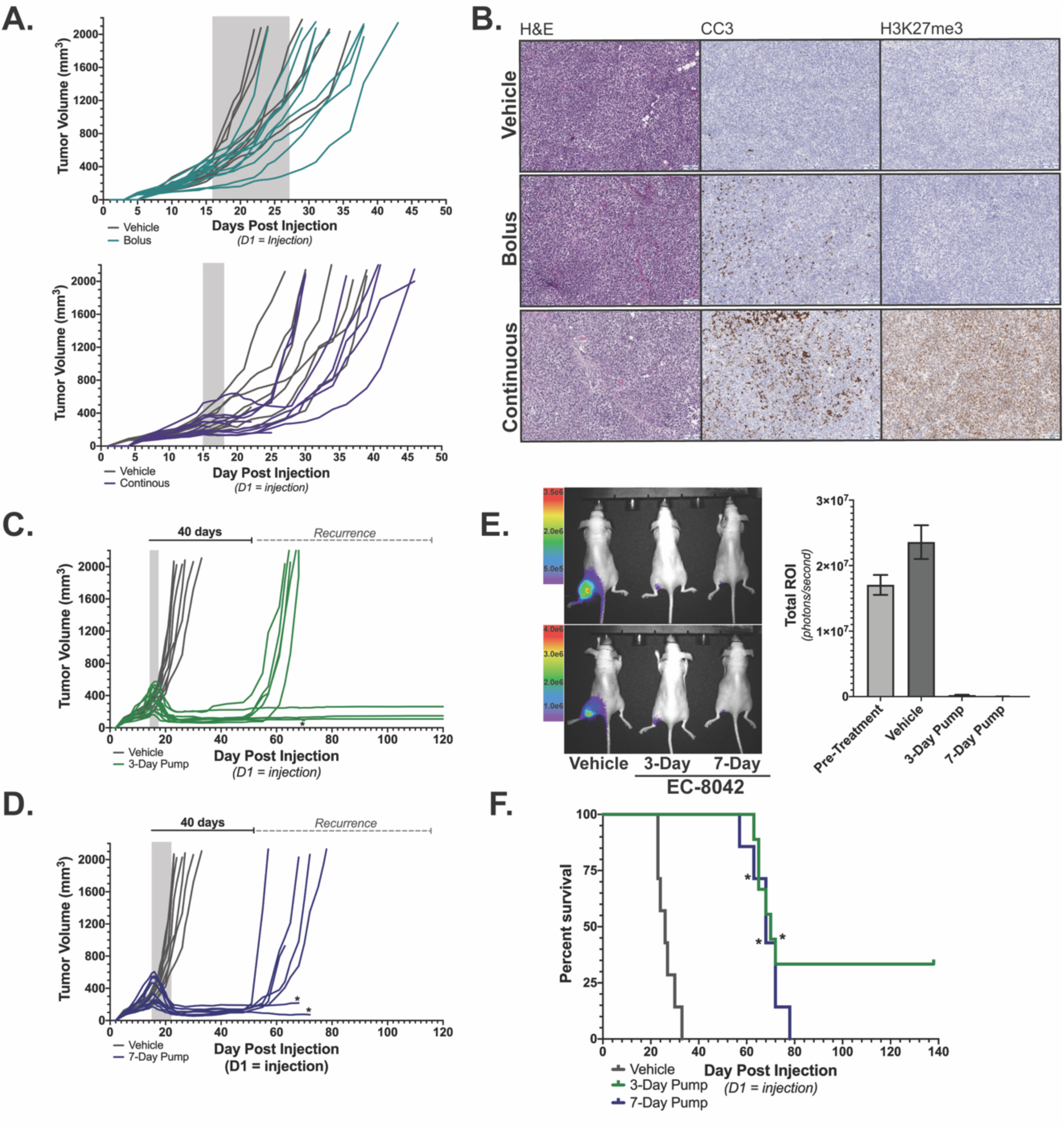
Continuous infusion of mithramycin shows activity in an intramuscular rhabdoid tumor xenograft model. **A:** Top, Spaghetti plot showing tumor volumes of individual tumors in mice bearing G401 xenografts treated with 1 mg/kg/dose mithramycin intraperitoneal M, W, F for two weeks (green) relative to vehicle control (gray). Bottom, Spaghetti plot showing tumor volumes of individual tumors in mice bearing G401 xenografts treated with 2.4 mg/kg of mithramycin (purple) or vehicle control (gray) administered continuously intraperitoneal over 72h. Most mice experienced a suppression or regression of tumor volume that persisted for more than 2 weeks following treatment. The shaded box indicates the duration of treatment. **B:** Immunohistochemistry analysis recapitulates the biochemistry described in vitro. G401 tumor sections at 10X magnification stained for cleaved caspase 3 (CC3; apoptosis) or H3K27me3. A marked increase in CC3 correlates with H3K27me3 staining is seen only in mice treated with the continuous infusion schedule but not bolus dosing. **C,D:** Prolonged durable response and cure of mice bearing G401 xenografts treated with **(C)** 30mg/kg EC8042 administered continuously over 72-hours or **(D)** 50 mg/kg EC8042 administered continuously over 144-hours. Treatment duration indicated by gray shaded box. Asterisks indicate an animal sacrificed due to unknown causes not related to tumor progression or drug toxicity (see text). **E:** Bioluminescence imaging of G401 rhabdoid tumor xenografts correlates with caliper measurements in 7A and 7B. Two mice per treatment group were imaged (left) and quantified in the bar graph (right). Error bars represent mean with SD. **F:** Kaplan-Meier survival curves indicating extended survival for mice bearing established G401 xenografts treated with the 3-day or 7-day continuous infusions of EC8042 in (C) and (D). Asterisks indicate an animal sacrificed due to unknown causes not related to tumor progression or drug toxicity (see text).

Importantly, continuous infusion recapitulated the mechanism of growth suppression described in vitro. There was a marked increase in H3K27me3 staining that correlated with apoptosis as measured by cleaved caspase 3 staining (Figure 5B). It is notable that this correlation was observed in vitro. Unfortunately, we could not escalate the dose further because of toxicity.

### EC8042 leads to marked tumor regression and mesenchymal differentiation in vivo

In an effort to improve the activity of mithramycin and increase the clinical relevance of the described effects, we evaluated the ability of the second-generation mithramycin analog EC8042 to recapitulate these effects. EC8042 is a second generation mithramycin analogue that is more than 10 times less toxic than mithramycin in multiple species but retains the hypersensitivity to SWI/SNF mutant tumors described above (15). Rhabdoid tumor cells were sensitive to EC8042 as mithramycin with a slightly higher IC50 and retained the described mesenchymal differentiation phenotype in vitro (Figure S6A, S6B). Mice with established 100 mm^3^ G401 rhabdoid tumor xenografts were treated with 3-days or 7-days of continuous infusion of EC8042 for a total dose of 30mg/kg or 50mg/kg, respectively. All mice in both cohorts experienced striking regressions of their well-established tumors including many with large tumors >400 mm^3^ (Figure 5C, 5D). Bioluminescent imaging in a subset of mice showed almost no detectable tumor at the end of infusion (Figure 5E). Retreatment was not possible due to IACUC limitations on a second pump implantation surgery. However, several mice were cured of their disease with a single 3-day infusion of EC8042 and never showed tumor recurrence 140 days after treatment (Figure 5F). The effects occurred with almost no observed toxicity as mice experienced minimal transient weight loss that resolved with cessation of drug infusion (Figure S6C).

Importantly, EC8042 drove the combined apoptotic and differentiation phenotype by the described mechanism of action. There was a global increase in H3K27me3 and reduction of H3K27ac (Figure 6A, 6B, S7A). The intense staining of H3K27me3 was not present in day 8, but there were no identifiable residual tumor cells (Figure 6C, S7A). Amplification of H3K27me3 positively correlated with cleaved caspase3 and reduction of Ki67 indicating suppression of proliferation and induction of apoptosis (Figure 6D, S7A, S7B). In addition, there was striking evidence of mesenchymal differentiation into bone with the appearance of trabecular-like ossification as well as cartilage and adipocytes (Figure 6E). The presence of osteoblasts and embedded osteocytes in the trabecular architecture provides further support for EC8042 inducing osteogenesis (Figure S7C). Finally, we confirmed calcification of the tumor tissue with microcomputed tomography (micro-CT) (Figure 6E). Importantly, the appearance of a mesenchymal differentiation phenotype is supported by rhabdoid tumor cell of origin studies indicating a mesenchymal origin and clinical evidence supporting mesenchymal features in patients (45). In addition, the PCA of BT12 cells treated with mithramycin clusters with gene signatures from bone. It is notable that Wnt3 (a known driver of osteogenesis) was one of the top promoters remodeled after mithramycin treatment (Figure S7D, data not shown). The differentiation phenotype is not fully penetrant and is represented by mixed lineage, likely due to transcriptional and epigenetic heterogeneity in the xenograft as well as influences of the micro-environment. Nevertheless, this phenotype leads to a significant increase in survival of the rhabdoid tumor xenografts treated with EC8042.

**Figure 6:**
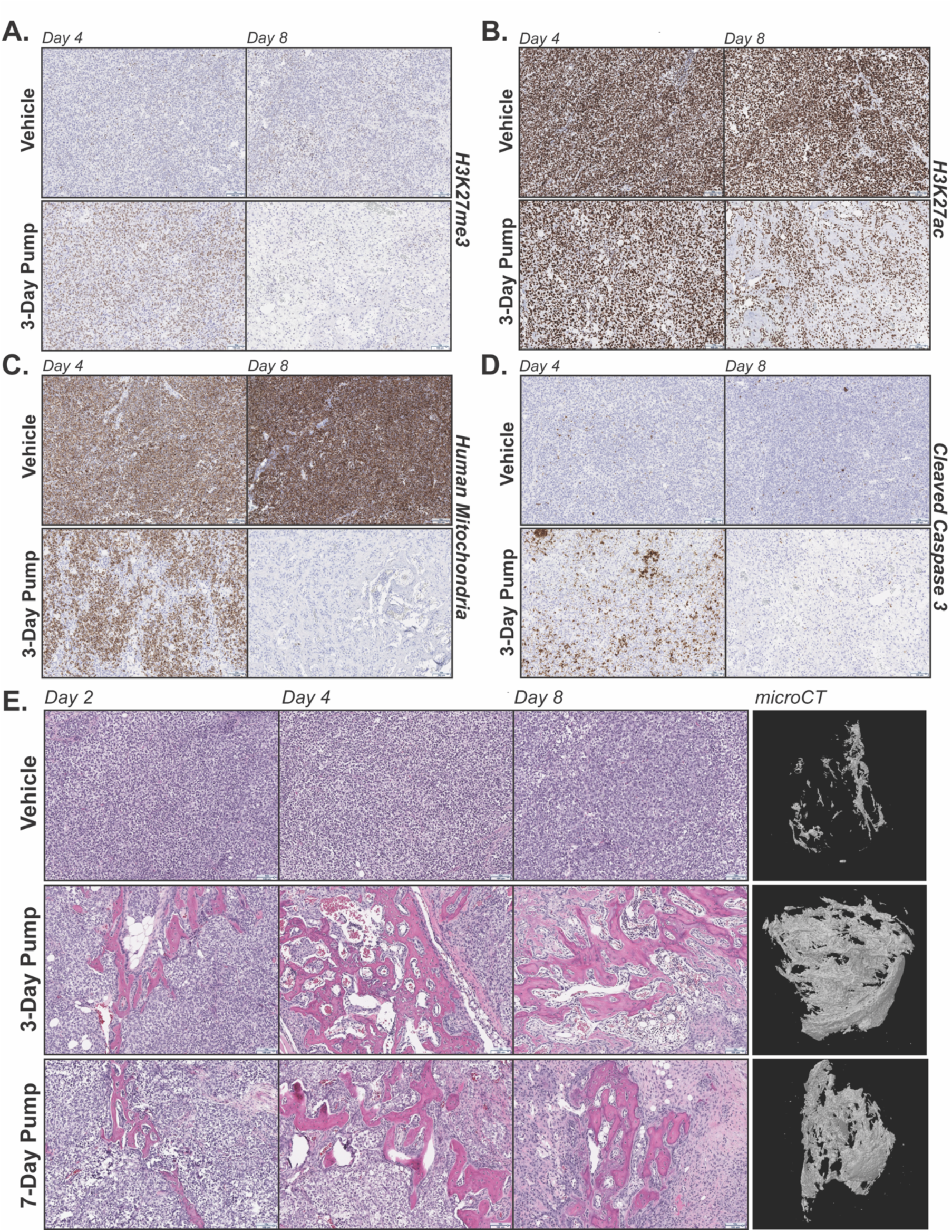
EC8042 induces a mixed phenotype that favors mesenchymal differentiation over apoptosis. **A-D:** 10X image of section of G401 treated tumors collected on day 4 (left) and day 8 (right) and stained for H3K27me3 **(A)**, H3K27ac **(B)**, human mitochondria **(C)**, and cleaved caspase 3 **(D)**. The sections compare vehicle to treatment started on day 1 with 30 mg/kg of EC8042 administered continuously for 72-hours (3-day pump). H3K27me3 increases and correlates with apoptosis (CC3) while H3K27ac decreases over time. H3K27ac and human mitochondrial staining decrease by day 4 compared with vehicle. Positive staining of H3K27me3 and CC3 is gone by day 8, as is human mitochondrial staining indicating no residual tumor. **E:** Immunohistochemistry analysis of H&E stains from G401 xenograft tumors on 1, 3, and 7-days after treatment with vehicle, 3-day EC8042 pump or 7-day EC8042 pump. EC8042 treated xenograft tumors exhibit evidence of mesenchymal differentiation compared to vehicle. microCT analysis of xenograft tumors on 7-days after treatment exhibit enhanced calcification compared to vehicle.

## DISCUSSION

In this study, we identify mithramycin as a direct inhibitor of the SWI/SNF chromatin remodeling complex. We show eviction of SWI/SNF from chromatin and the induction of a striking cellular response characterized by chromatin compartment remodeling, a change in the distribution of H3K27ac and differential changes in chromatin accessibility that drive both an apoptotic and differentiation phenotype. This genotype and phenotype is clearly captured in vitro in modeling experiments and in a multi-omic analysis that links gene expression changes to alterations in chromatin structure and accessibility. Importantly, the epigenetic reprogramming favors regions near the transcriptional start site and not long-range enhancers. Together, these effects explain the striking sensitivity of this tumor to this drug and provides a new therapeutic option for these notoriously chemo-refractory patients, particularly with the less-toxic analogue EC8042. Importantly, EC8042 emerges as a targeted agent for two different driver oncogenes in two different tumors, Ewing sarcoma and rhabdoid tumor. It is likely that this response to the release of the aberrant SWI/SNF in both tumor types accounts for the heightened sensitivity of these tumors to these compounds. It is tempting to speculate that these compounds may be active in the 20% of tumors characterized by mutated SWI/SNF as suggested by the in vitro screening data reported in this manuscript. This increases the attractiveness of this agent as a clinical candidate due to the rarity of these tumors.

Importantly, in addition to a new therapeutic option, this study provides important insight into the biology of RT. It has been suggested that the loss of SMARCB1 leads to a SWI/SNF complex that binds to chromatin with lower affinity and is distributed aberrantly throughout the genome. Consistent with this idea, we identified a small molecule that inhibits oncogenic SWI/SNF with some selectivity relative to wild-type. We show that eviction of SWI/SNF only occurs in RT cells and not in SWI/SNF wild-type U2OS cells. In addition, the toxicity profile, particularly for EC8042, demonstrates a favorable therapeutic window. This suggests effects on the tumor cells but not on the normal cells which are known to possess wild-type SWI/SNF.

Interestingly, in this study we show a striking schedule dependence of these tumors for both mithramycin and EC8042. We demonstrate that both compounds are more effective as a continuous infusion that is able to induce both an apoptotic and a differentiation endpoint of therapy. With mithramycin, this endpoint is exposure dependent where transient high concentration exposures favor apoptosis while low concentrations over time favor differentiation. It is an important observation that the high-concentration, transient exposure that drives the apoptotic phenotype is less effective in the xenografts than the continuous infusion. Only the continuous infusion induces durable responses and even complete cures and importantly drives the described mechanism of action with cellular differentiation, increased H3K27me3 and loss of H3K27ac observed in the tissue. Notably, in the case where differentiation agents have been identified in a number of tumors, these tend to be extremely active in patients with some examples being arsenic and ATRA for APL, trabectedin for myxoid liposarcoma, and retinoids for neuroblastoma (46–48). Furthermore, these observations provide a biomarker of activity. The induction of H3K27me3 and loss of H3K27ac staining of the FFPE tumor tissue was quite striking and only occurred in tumors that responded to the drug.

Finally, this study provides important insight into the mechanism of mithramycin. Mithramycin was originally identified as an anti-cancer agent in the 1950’s. While it showed some activity in the clinic, it fell out of favor due to its narrow toxicity profile. It has always been referred to as an SP1 inhibitor with recent activity suggesting activity against ETS transcription factors and EWS-FLI1, the oncogenic driver of Ewing sarcoma (13). The data in this study suggests that, at least in these cells, the activity against SP1 is the result of SWI/SNF blockade and not the primary mechanism of action. This allows the identification of more effective analogues, such as EC8042, for Ewing sarcoma and other tumors that are sensitive to this class of compounds. In addition, in tumors dependent on SP1, it provides insight into approaches to develop novel combination therapies that perhaps amplify the targeting of SP1.

Together these data provide important insight into the biology and therapeutic targeting of SWI/SNF, a novel therapeutic option for the notoriously chemo-refractory rhabdoid tumor and mechanistic insight into mithramycin analogs that has the potential to impact a broad range of cancers characterized by dysregulation of SWI/SNF.

## ACKNOWLEDGEMENTS

Research reported in this manuscript was supported by the National Cancer Institute of the National Institutes of Health under the award number F31-CA236300-01 (M.H.C.) and internal funds from the Van Andel Research Institute (T.J.T. and P.J.G.). The authors would like to thank the Van Andel Genomics, Bioinformatics and Biostatistics, and Pathology and Biorepository Cores for providing next generation sequencing facilities, formal analysis and immunohistochemistry analysis. The authors would like to thank EntreChem for the use of EC8042 and Robert Vaughan from the Rothbart lab for technical advice. Finally, we would like to thank the patients, especially J.S. and family.

## AUTHOR CONTRIBUTION

Conceptualization, M.H.C., and P.J.G.; Methodology, M.H.C., B.K.J., E.A.B., T.J.T., B.O.W., P.J.G.; Investigation, M.H.C., E.A.B., K.M.S., S.M.G.; Formal Analysis: M.H.C., B.K.J., L.H., I.B., Z.B.M., G.E.F., B.O.W., T.J.T., P.J.G.; Writing, M.H.C., B.K.J., P.J.G.; Funding Acquisition, M.H.C., T.J.T., P.J.G.; Resources, P.J.G.; Supervision, B.O.W., T.J.T., P.J.G.

## REFERENCES

1. Versteege I, Sevenet N, Lange J, Rousseau-Merck MF, Ambros P, Handgretinger R, et al. Truncating mutations of hSNF5/INI1 in aggressive paediatric cancer. Nature 1998;394(6689):203–6 doi 10.1038/28212.

2. Brennan B, Stiller C, Bourdeaut F. Extracranial rhabdoid tumours: what we have learned so far and future directions. The Lancet Oncology 2013;14(8):e329–36 doi 10.1016/s1470-2045(13)70088-3.

3. Ginn KF, Gajjar A. Atypical teratoid rhabdoid tumor: current therapy and future directions. Frontiers in oncology 2012;2:114 doi 10.3389/fonc.2012.00114.

4. Nakayama RT, Pulice JL, Valencia AM, McBride MJ, McKenzie ZM, Gillespie MA, et al. SMARCB1 is required for widespread BAF complex-mediated activation of enhancers and bivalent promoters. Nature genetics 2017;49(11):1613–23 doi 10.1038/ng.3958.

5. Wang X, Lee RS, Alver BH, Haswell JR, Wang S, Mieczkowski J, et al. SMARCB1-mediated SWI/SNF complex function is essential for enhancer regulation. Nature genetics 2017;49(2):289–95 doi 10.1038/ng.3746.

6. Erkek S, Johann PD, Finetti MA, Drosos Y, Chou HC, Zapatka M, et al. Comprehensive Analysis of Chromatin States in Atypical Teratoid/Rhabdoid Tumor Identifies Diverging Roles for SWI/SNF and Polycomb in Gene Regulation. Cancer cell 2019;35(1):95–110.e8 doi 10.1016/j.ccell.2018.11.014.

7. Wang X, Sansam CG, Thom CS, Metzger D, Evans JA, Nguyen PT, et al. Oncogenesis caused by loss of the SNF5 tumor suppressor is dependent on activity of BRG1, the ATPase of the SWI/SNF chromatin remodeling complex. Cancer research 2009;69(20):8094–101 doi 10.1158/0008-5472.Can-09-0733.

8. Wang X, Wang S, Troisi EC, Howard TP, Haswell JR, Wolf BK, et al. BRD9 defines a SWI/SNF sub-complex and constitutes a specific vulnerability in malignant rhabdoid tumors. Nature communications 2019;10(1):1881 doi 10.1038/s41467-019-09891-7.

9. Michel BC, D’Avino AR, Cassel SH, Mashtalir N, McKenzie ZM, McBride MJ, et al. A non-canonical SWI/SNF complex is a synthetic lethal target in cancers driven by BAF complex perturbation. Nature cell biology 2018;20(12):1410–20 doi 10.1038/s41556-018-0221-1.

10. Wilson BG, Wang X, Shen X, McKenna ES, Lemieux ME, Cho YJ, et al. Epigenetic antagonism between polycomb and SWI/SNF complexes during oncogenic transformation. Cancer cell 2010;18(4):316–28 doi 10.1016/j.ccr.2010.09.006.

11. Kia SK, Gorski MM, Giannakopoulos S, Verrijzer CP. SWI/SNF mediates polycomb eviction and epigenetic reprogramming of the INK4b-ARF-INK4a locus. Molecular and cellular biology 2008;28(10):3457–64 doi 10.1128/mcb.02019-07.

12. Knutson SK, Warholic NM, Wigle TJ, Klaus CR, Allain CJ, Raimondi A, et al. Durable tumor regression in genetically altered malignant rhabdoid tumors by inhibition of methyltransferase EZH2. Proceedings of the National Academy of Sciences of the United States of America 2013;110(19):7922–7 doi 10.1073/pnas.1303800110.

13. Grohar PJ, Woldemichael GM, Griffin LB, Mendoza A, Chen QR, Yeung C, et al. Identification of an inhibitor of the EWS-FLI1 oncogenic transcription factor by high-throughput screening. Journal of the National Cancer Institute 2011;103(12):962–78 doi 10.1093/jnci/djr156.

14. Boulay G, Sandoval GJ, Riggi N, Iyer S, Buisson R, Naigles B, et al. Cancer-Specific Retargeting of BAF Complexes by a Prion-like Domain. Cell 2017;171(1):163–78.e19 doi 10.1016/j.cell.2017.07.036.

15. Osgood CL, Maloney N, Kidd CG, Kitchen-Goosen S, Segars L, Gebregiorgis M, et al. Identification of Mithramycin Analogues with Improved Targeting of the EWS-FLI1 Transcription Factor. Clinical cancer research : an official journal of the American Association for Cancer Research 2016;22(16):4105–18 doi 10.1158/1078-0432.Ccr-15-2624.

16. Harlow ML, Chasse MH, Boguslawski EA, Sorensen KM, Gedminas JM, Kitchen-Goosen SM, et al. Trabectedin Inhibits EWS-FLI1 and Evicts SWI/SNF from Chromatin in a Schedule-dependent Manner. Clinical cancer research : an official journal of the American Association for Cancer Research 2019;25(11):3417–29 doi 10.1158/1078-0432.Ccr-18-3511.

17. Harlow ML, Maloney N, Roland J, Guillén-Navarro MJ, Easton MK, Kitchen-Goosen SM, et al. Lurbinectedin inactivates the Ewing sarcoma oncoprotein EWS-FLI1 by redistributing it within the nucleus. Cancer research 2016:canres.0568.2016 doi 10.1158/0008-5472.CAN-16-0568.

18. Dobin A, Davis CA, Schlesinger F, Drenkow J, Zaleski C, Jha S, et al. STAR: ultrafast universal RNA-seq aligner. Bioinformatics (Oxford, England) 2013;29(1):15–21 doi 10.1093/bioinformatics/bts635.

19. Robinson MD, McCarthy DJ, Smyth GK. edgeR: a Bioconductor package for differential expression analysis of digital gene expression data. Bioinformatics (Oxford, England) 2010;26(1):139–40 doi 10.1093/bioinformatics/btp616.

20. Robinson MD, Oshlack A. A scaling normalization method for differential expression analysis of RNA-seq data. Genome biology 2010;11(3):R25 doi 10.1186/gb-2010-11-3-r25.

21. Ritchie ME, Phipson B, Wu D, Hu Y, Law CW, Shi W, et al. limma powers differential expression analyses for RNA-sequencing and microarray studies. Nucleic acids research 2015;43(7):e47 doi 10.1093/nar/gkv007.

22. Law CW, Chen Y, Shi W, Smyth GK. voom: precision weights unlock linear model analysis tools for RNA-seq read counts. Genome biology 2014;15(2):R29 doi 10.1186/gb-2014-15-2-r29.

23. Sergushichev AA. An algorithm for fast preranked gene set enrichment analysis using cumulative statistic calculation. bioRxiv 2016:060012 doi 10.1101/060012.

24. Yu G, He QY. ReactomePA: an R/Bioconductor package for reactome pathway analysis and visualization. Molecular bioSystems 2016;12(2):477–9 doi 10.1039/c5mb00663e.

25. Rojas-Pena ML, Olivares-Navarrete R, Hyzy S, Arafat D, Schwartz Z, Boyan BD, et al. Characterization of distinct classes of differential gene expression in osteoblast cultures from non-syndromic craniosynostosis bone. Journal of genomics 2014;2:121–30 doi 10.7150/jgen.8833.

26. Faust GG, Hall IM. SAMBLASTER: fast duplicate marking and structural variant read extraction. Bioinformatics (Oxford, England) 2014;30(17):2503–5 doi 10.1093/bioinformatics/btu314.

27. Li H, Handsaker B, Wysoker A, Fennell T, Ruan J, Homer N, et al. The Sequence Alignment/Map format and SAMtools. Bioinformatics (Oxford, England) 2009;25(16):2078–9 doi 10.1093/bioinformatics/btp352.

28. Lun AT, Smyth GK. csaw: a Bioconductor package for differential binding analysis of ChIP-seq data using sliding windows. Nucleic acids research 2016;44(5):e45 doi 10.1093/nar/gkv1191.

29. Lun AT, Smyth GK. De novo detection of differentially bound regions for ChIP-seq data using peaks and windows: controlling error rates correctly. Nucleic acids research 2014;42(11):e95 doi 10.1093/nar/gku351.

30. Amemiya HM, Kundaje A, Boyle AP. The ENCODE Blacklist: Identification of Problematic Regions of the Genome. Scientific Reports 2019;9(1):9354 doi 10.1038/s41598-019-45839-z.

31. Corces MR, Trevino AE, Hamilton EG, Greenside PG, Sinnott-Armstrong NA, Vesuna S, et al. An improved ATAC-seq protocol reduces background and enables interrogation of frozen tissues. Nature methods 2017;14(10):959–62 doi 10.1038/nmeth.4396.

32. Risso D, Ngai J, Speed TP, Dudoit S. Normalization of RNA-seq data using factor analysis of control genes or samples. Nature Biotechnology 2014;32:896 doi 10.1038/nbt.2931 https://www.nature.com/articles/nbt.2931#supplementary-information.

33. Risso D, Schwartz K, Sherlock G, Dudoit S. GC-Content Normalization for RNA-Seq Data. BMC Bioinformatics 2011;12(1):480 doi 10.1186/1471-2105-12-480.

34. Deng C, Daley T, Smith AD. Applications of species accumulation curves in large-scale biological data analysis. Quantitative biology (Beijing, China) 2015;3(3):135–44 doi 10.1007/s40484-015-0049-7.

35. Chun HE, Lim EL, Heravi-Moussavi A, Saberi S, Mungall KL, Bilenky M, et al. Genome-Wide Profiles of Extra-cranial Malignant Rhabdoid Tumors Reveal Heterogeneity and Dysregulated Developmental Pathways. Cancer cell 2016;29(3):394–406 doi 10.1016/j.ccell.2016.02.009.

36. Ernst J, Kellis M. ChromHMM: automating chromatin-state discovery and characterization. Nature methods 2012;9:215 doi 10.1038/nmeth.1906 https://www.nature.com/articles/nmeth.1906#supplementary-information.

37. Ernst J, Kellis M. Chromatin-state discovery and genome annotation with ChromHMM. Nature Protocols 2017;12:2478 doi 10.1038/nprot.2017.124.

38. Roadmap Epigenomics C, Kundaje A, Meuleman W, Ernst J, Bilenky M, Yen A, et al. Integrative analysis of 111 reference human epigenomes. Nature 2015;518:317 doi 10.1038/nature14248 https://www.nature.com/articles/nature14248#supplementary-information.

39. Teicher BA, Polley E, Kunkel M, Evans D, Silvers T, Delosh R, et al. Sarcoma Cell Line Screen of Oncology Drugs and Investigational Agents Identifies Patterns Associated with Gene and microRNA Expression. Molecular cancer therapeutics 2015;14(11):2452–62 doi 10.1158/1535-7163.Mct-15-0074.

40. Quinn J, Fyrberg AM, Ganster RW, Schmidt MC, Peterson CL. DNA-binding properties of the yeast SWI/SNF complex. Nature 1996;379(6568):844–7 doi 10.1038/379844a0.

41. Sastry M, Patel DJ. Solution structure of the mithramycin dimer-DNA complex. Biochemistry 1993;32(26):6588–604 doi 10.1021/bi00077a012.

42. Snyder RC, Ray R, Blume S, Miller DM. Mithramycin blocks transcriptional initiation of the c-myc P1 and P2 promoters. Biochemistry 1991;30(17):4290–7 doi 10.1021/bi00231a027.

43. Remsing LL, Bahadori HR, Carbone GM, McGuffie EM, Catapano CV, Rohr J. Inhibition of c-src transcription by mithramycin: structure-activity relationships of biosynthetically produced mithramycin analogues using the c-src promoter as target. Biochemistry 2003;42(27):8313–24 doi 10.1021/bi034091z.

44. Weintraub AS, Li CH, Zamudio AV, Sigova AA, Hannett NM, Day DS, et al. YY1 Is a Structural Regulator of Enhancer-Promoter Loops. Cell 2017;171(7):1573–88.e28 doi 10.1016/j.cell.2017.11.008.

45. Rorke LB, Packer RJ, Biegel JA. Central nervous system atypical teratoid/rhabdoid tumors of infancy and childhood: definition of an entity. Journal of neurosurgery 1996;85(1):56–65 doi 10.3171/jns.1996.85.1.0056.

46. Forni C, Minuzzo M, Virdis E, Tamborini E, Simone M, Tavecchio M, et al. Trabectedin (ET-743) promotes differentiation in myxoid liposarcoma tumors. Molecular cancer therapeutics 2009;8(2):449–57 doi 10.1158/1535-7163.Mct-08-0848.

47. Sidell N, Altman A, Haussler MR, Seeger RC. Effects of retinoic acid (RA) on the growth and phenotypic expression of several human neuroblastoma cell lines. Experimental cell research 1983;148(1):21–30 doi 10.1016/0014-4827(83)90184-2.

48. Flynn PJ, Miller WJ, Weisdorf DJ, Arthur DC, Brunning R, Branda RF. Retinoic acid treatment of acute promyelocytic leukemia: in vitro and in vivo observations. Blood 1983;62(6):1211–7.

49. Tate JG, Bamford S, Jubb HC, Sondka Z, Beare DM, Bindal N, et al. COSMIC: the Catalogue Of Somatic Mutations In Cancer. Nucleic acids research 2019;47(D1):D941–d7 doi 10.1093/nar/gky1015.

