## Supplemental Material for "Direct therapeutic targeting of SWI/SNF induces epigenetic reprogramming and durable tumor regression in rhabdoid tumor"

### SUPPLEMENTAL INFORMATION

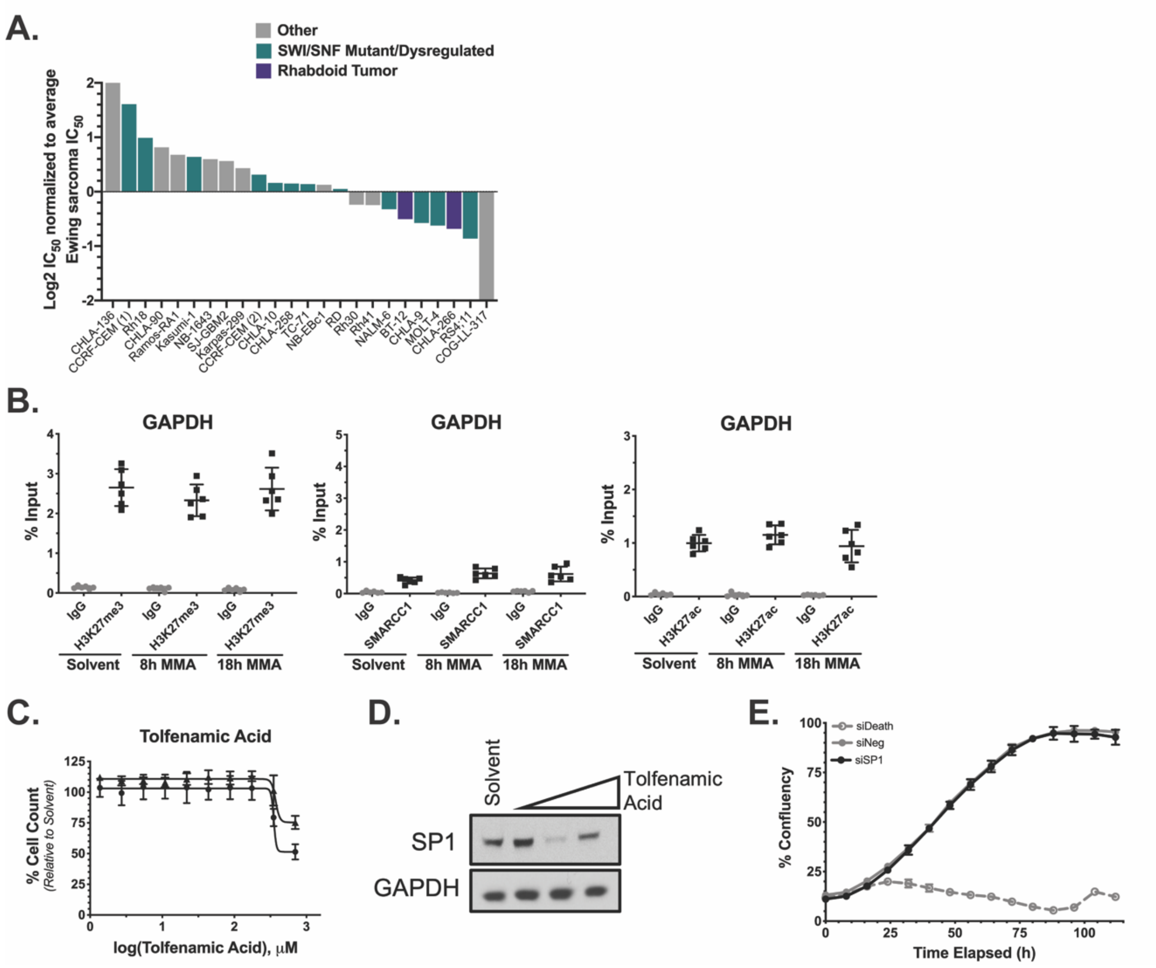


**Supplementary Figure 1: Mithramycin dependent SWI/SNF eviction is not due to SP1 activity and does not induce reprogramming at housekeeping genes**

**A:** A pediatric cell line viability screen with mithramycin. Cell lines with mutated or dysregulated SWI/SNF cluster towards the right on the graph indicating these cell lines are more sensitive to mithramycin. Data analyzed from *(16)*.
**B:** Chromatin immunoprecipitation of IgG, SMARCC1 (left), H3K27ac (middle) or H3K27me3 (right) at the control locus, *GAPDH*.
**C:** Knockdown of SP1 (black) does not affect BT12 rhabdoid tumor cell proliferation compared with a siNeg (gray, solid line) negative control as measured by live cell imaging. siDeath (gray, dotted line) is a positive control for knockdown efficiency.
**D,E:** Dose response curve showing limited sensitivity of RT cells to tolfenamic acid treatment in BT12 (circle) and G401 (triangle) cells **(D)** despite suppression in SP1 expression as measured by western blot following exposure to 0.5µM, 1µM, and 2µM for 48-hours **(E).**

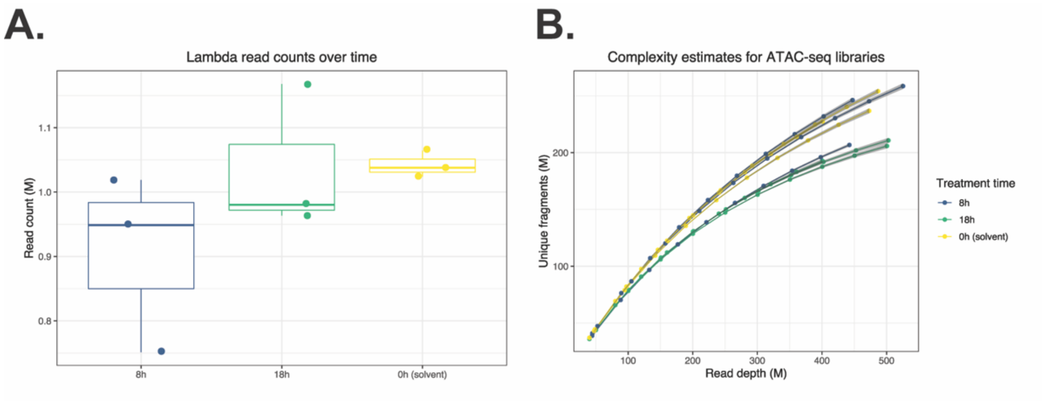


**Supplementary Figure 2: Lambda read depth and library complexity for ATAC sequencing**

**A:** Lambda spike-in reads that have been de-duplicated, stratified by replicate and mithramycin treatment over time that corresponds to RUV-based accessibility normalization for ATAC-seq libraries in Figures 3.
**B:** Estimated ATAC-seq library complexity of individual replicates up to 10x the initial library size (first data point shown), demonstrating fundamental differences in accessibility, and by extension, library complexity from mithramycin treatment, demanding spike-in normalization procedures for adequate comparisons across drug treatment as performed in Figure 3.

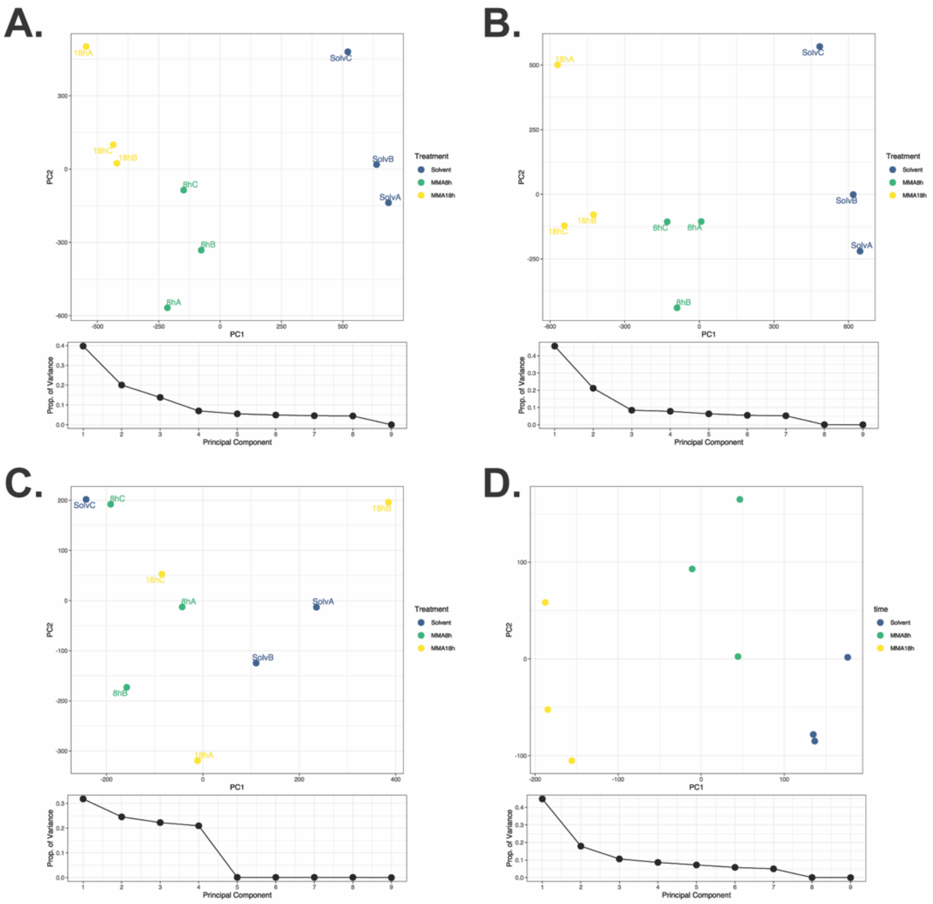


**Supplementary Figure 3: Lambda spike-in normalization for ATAC and ChIP sequencing**

**A:** Principal component analysis of ATAC-seq libraries prior to normalization with lambda spike-in controls. Only the first 2 principal components are shown with a scree plot showing all 9 components and variance explained.
**B:** Principal component analysis of ATAC-seq libraries post-normalization with lambda spike-in controls using a k of 1 for RUVg.
**C:** Principal component analysis of ATAC-seq libraries post-normalization with lambda spike-in controls using a k of 4 for RUVg. Notably, treatment level clusters are lost using the spike-ins as control “genes” within the first 4 factors, suggesting that library complexity can be accounted for or confounding if not addressed as dominant sources of variation in the data.
**D:** Principal component analysis of H3K27Ac ChIP-seq data following regressing principal component 1 out of the data.

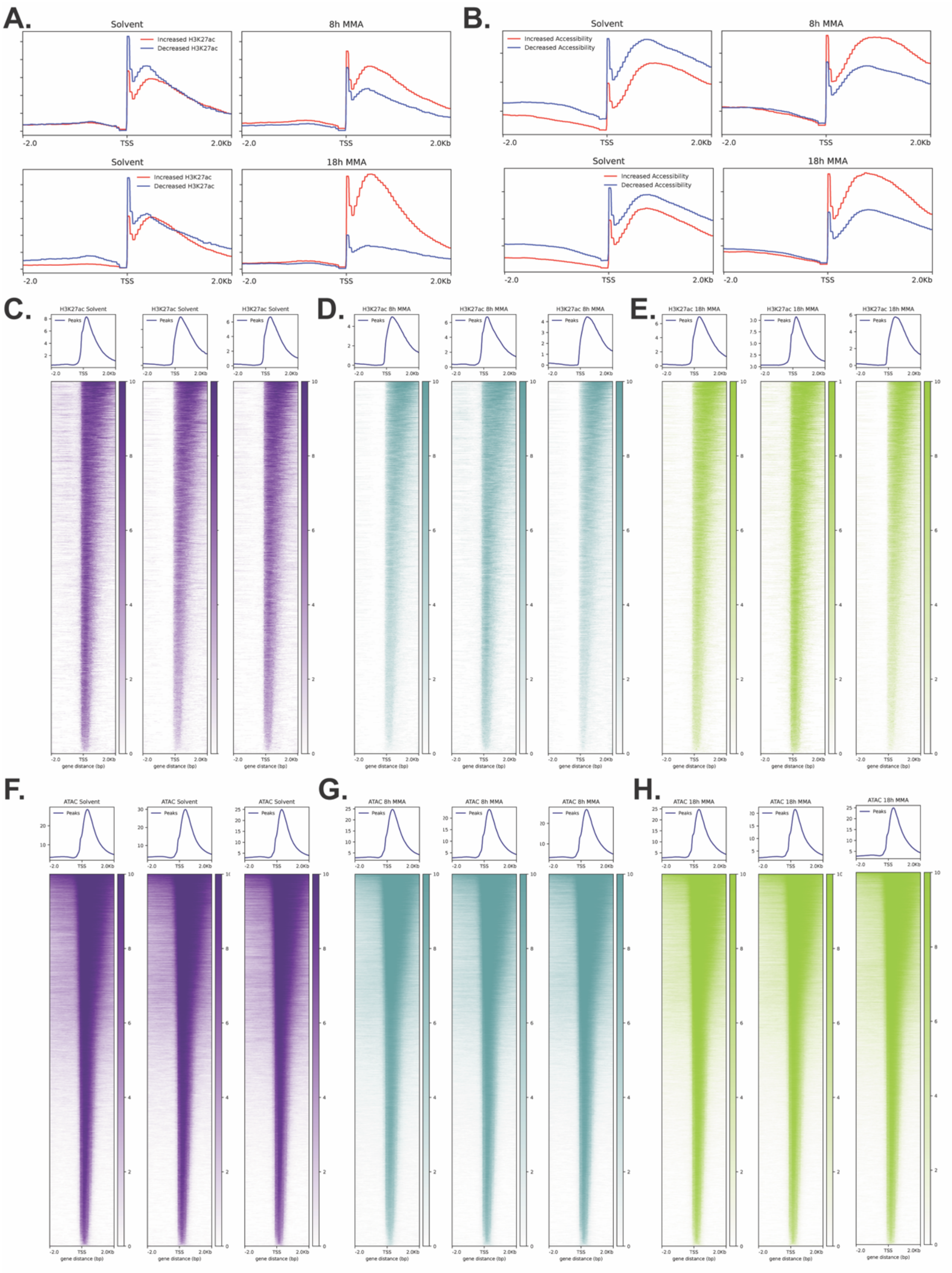


**Supplementary Figure 4: Genome-wide peak analysis for ATAC and ChIP sequencing**

**A:** Profile traces that correspond with the heatmaps plotted in Figure 3C for regions that increase in H3K27ac (red) and regions that decrease in H3K27ac (blue) from solvent to 8-hours (top) and 18-hours (bottom) mithramycin treatment.

**Supplemental Figure 4 Continued**

**B:** Profile traces that correspond with the heatmaps plotted in Figure 3C for regions that increase in accessibility (red) and regions that decrease in accessibility (blue) from solvent to 8-hours (top) and 18-hours (bottom) mithramycin treatment.
**C-E:** Heatmaps and profile tracing showing H3K27ac ChIP-seq peaks following mithramycin treatment for all three experimental replicates. A 2kb window is centered on the TSS.
**F-H:** Heatmaps and profile tracing showing ATAC-seq peaks following mithramycin treatment for all three experimental replicates. A 2kb window is centered on the TSS.

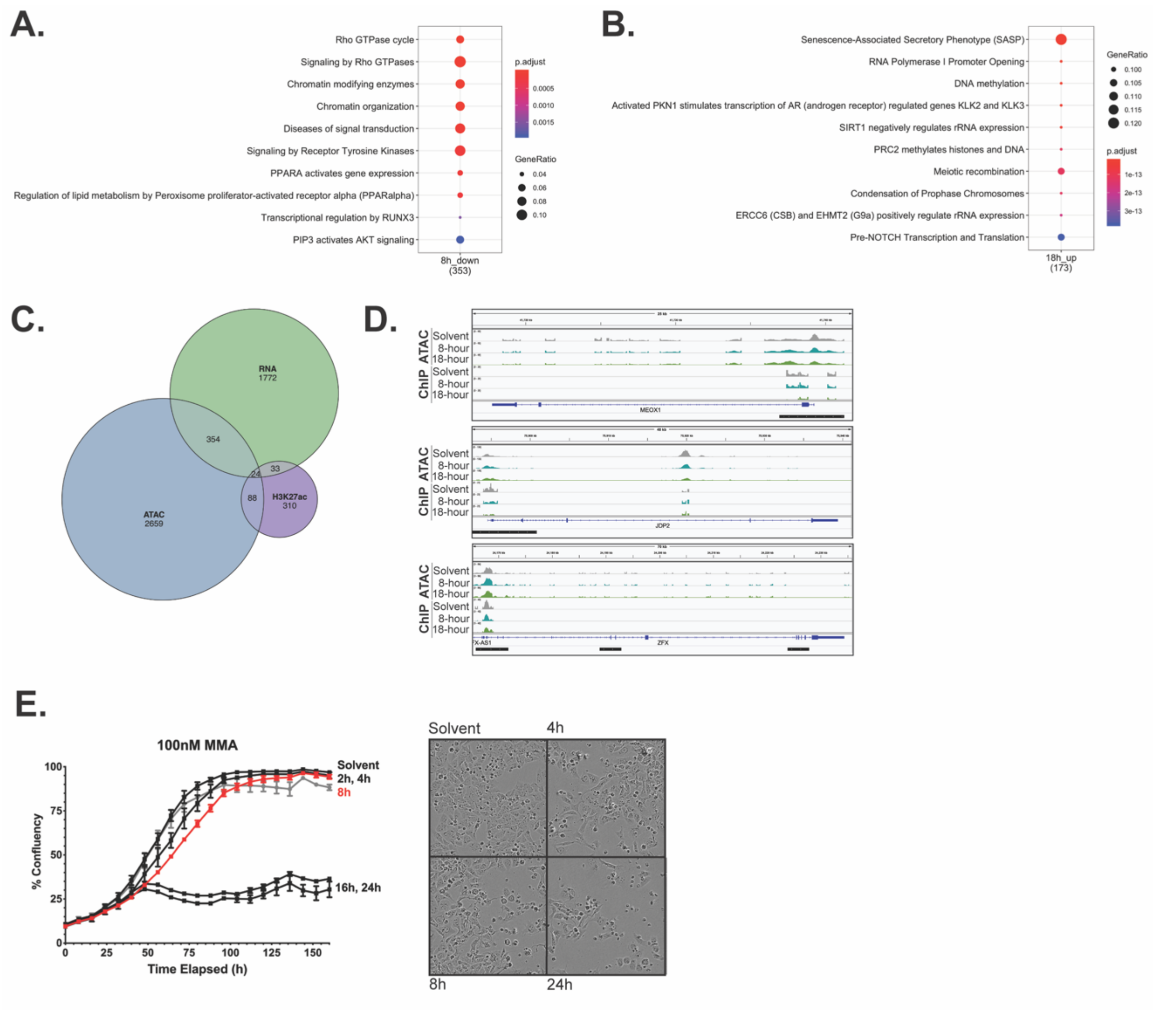


**Supplementary Figure 5: Genomic and cellular analyses of pathways up or down-regulated following mithramycin treatment**

**A, B:** RNA-seq reactome pathway analysis of the genes downregulated after 8-hours **(A)** and up-regulated after 18-hours **(B)** mithramycin treatment.
**C:** Venn diagram of overlapping gene sets with at least a 1log FC decrease in both 8-hours and 18-hours mithramycin treatment compared with solvent. 25 genes were down-regulated at both time points for RNA-seq, ATAC-seq, and H3K27ac ChIP-seq.
**D:** IGV tracks of three genes that were down-regulated in the multi-omic analysis. Black bars indicate promoters from hg19 within 2kb of the TSS.
**E:** Time course of 100nM mithramycin exposure in G401 cells. Cells were treated with 100nM MMA for the indicated times followed by a replacement of drug-free media (left). After 8-hours (red) of mithramycin exposures, the cells have an irreversible suppression of proliferation compared to solvent control. Live cell images of G401 cells treated with 100nM mithramycin at the indicated times (left).

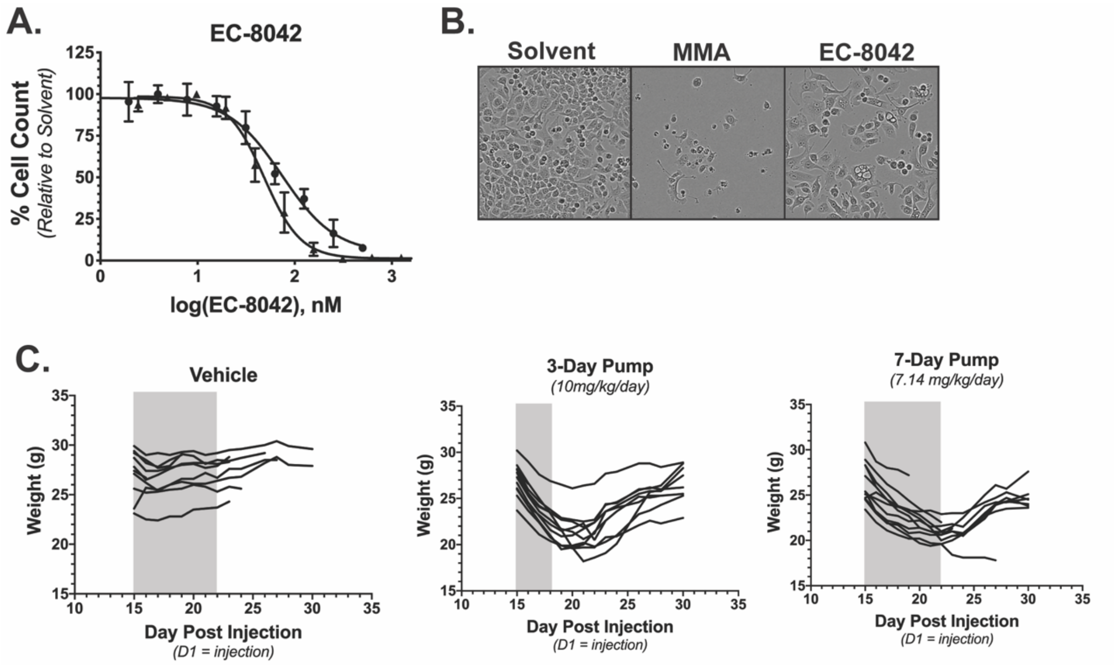


**Supplementary Figure 6: In vitro analysis and toxicity analysis of EC8042 treatment**

**A:** Dose response curve of BT12 (circle) and G401 (triangle) cells treated with EC8042. Both rhabdoid tumor cell lines are sensitive to EC8042.
**B:** Live cell images in BT12 cells treated with mithramycin and EC8042. EC8042 (200nM) leads to lipid accumulation and a flattened cell phenotype compared with mithramycin (100nM).
**C:** Mice treated with EC8042 have reversible body mass loss during treatment with the 3-day pump (middle) and the 7-day pump (right) compared to vehicle (left). However, body weight recovers once treatment ends.

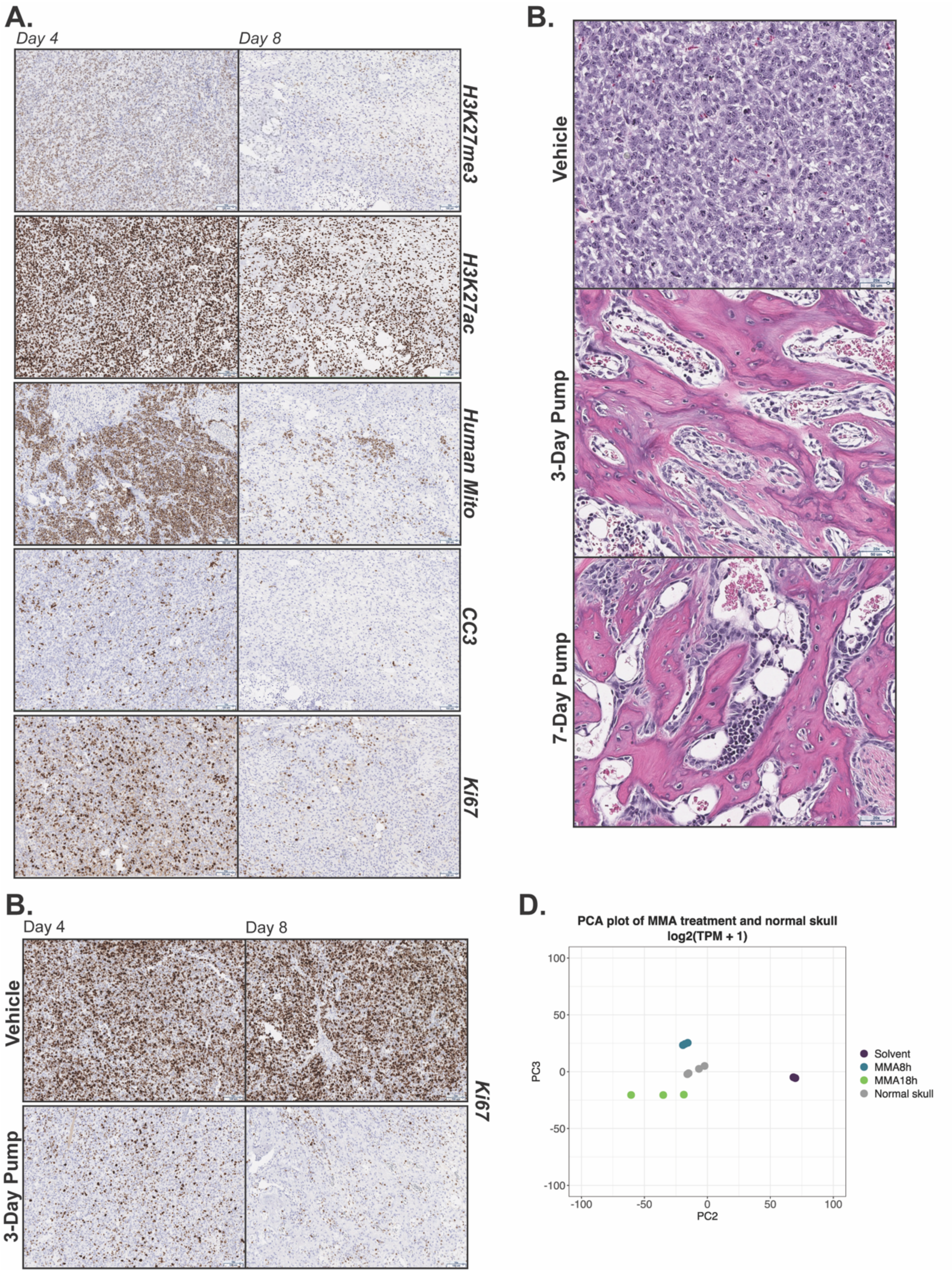


**Supplementary Figure 7: In vivo analysis of rhabdoid tumor xenografts following EC8042 treatment**

**Supplemental Figure 7 Continued**

**A:** Immunohistochemistry analysis of G401 xenograft tumors on 3 days (left, Day 4) and 7-days (right, Day 8) after treatment with vehicle or 7-day pump of EC8042. 10x magnification of H3K27me3, H3K27ac, human mitochondria, cleaved caspase 3, and Ki67 correlate with vehicle and 3-day EC8042 pump in Figure 6.
**B:** Immunohistochemistry analysis of G401 xenograft tumors on 3 days (left, Day 4) and 7-days (right, Day 8) after treatment with vehicle or 3-day pump of EC8042. 10x magnification of Ki67 correlates with the IHC shown in Figure 6.
**C:** Immunohistochemistry analysis of G401 xenograft tumors on 7 days (day 8) after treatment with vehicle, 3-day pump or 7-day pump of EC8042. 20x magnification of H&E shows osteoblasts and imbedded osteocytes in the trabecular architecture of treated xenograft tissue.
**D:** PCA analysis of mithramycin-treated BT12 cells with normal skull. Mithramycin treated cells cluster more with skull compared with solvent.
